# Non-parametric polygenic risk prediction using partitioned GWAS summary statistics

**DOI:** 10.1101/370064

**Authors:** Sung Chun, Maxim Imakaev, Daniel Hui, Nikolaos A. Patsopoulos, Benjamin M. Neale, Sekar Kathiresan, Nathan O. Stitziel, Shamil R. Sunyaev

**Affiliations:** Division of Genetics, Brigham and Women’s Hospital, Boston, Massachusetts, 02115, USA; Department of Biomedical Informatics, Harvard Medical School, Boston, Massachusetts, 02115, USA; Broad Institute of Harvard and MIT, Cambridge, Massachusetts, 02142, USA; Altius Institute for Biomedical Sciences, Seattle, Washington, 98121, USA; Systems Biology and Computer Science Program, Ann Romney Center for Neurological Diseases, Department of Neurology, Brigham & Women’s Hospital, Boston, 02115 MA, USA; Analytic and Translational Genetics Unit, Massachusetts General Hospital, Boston, Massachusetts, 02114, USA; Center for Human Genetic Research, Massachusetts General Hospital, Boston, Massachusetts, 02114, USA; Cardiovascular Research Center, Massachusetts General Hospital, Boston, Massachusetts, 02114, USA; Cardiovascular Division, Department of Medicine, Washington University School of Medicine, Saint Louis, Missouri, 63110, USA; Department of Genetics, Washington University School of Medicine, Saint Louis, Missouri, 63110, USA; McDonnell Genome Institute, Washington University School of Medicine, Saint Louis, Missouri, 63110, USA

**Author notes:** These authors contributed equally to this work (Sung Chun, Maxim Imakaev). Correspondence (NOS). Correspondence (SRS).

## Abstract

In complex trait genetics, the ability to predict phenotype from genotype is the ultimate measure of our understanding of genetic architecture underlying the heritability of a trait. A complete understanding of the genetic basis of a trait should allow for predictive methods with accuracies approaching the trait’s heritability. The highly polygenic nature of quantitative traits and most common phenotypes has motivated the development of statistical strategies focused on combining myriad individually non-significant genetic effects. Now that predictive accuracies are improving, there is a growing interest in practical utility of such methods for predicting risk of common diseases responsive to early therapeutic intervention. However, existing methods require individual level genotypes or depend on accurately specifying the genetic architecture underlying each disease to be predicted. Here, we propose a polygenic risk prediction method that does not require explicitly modeling any underlying genetic architecture. We start with summary statistics in the form of SNP effect sizes from a large GWAS cohort. We then remove the correlation structure across summary statistics arising due to linkage disequilibrium and apply a piecewise linear interpolation on conditional mean effects. In both simulated and real datasets, this new non-parametric shrinkage (NPS) method can reliably allow for linkage disequilibrium in summary statistics of 5 million dense genome-wide markers and consistently improves prediction accuracy. We show that NPS improves the identification of groups at high risk for Breast Cancer, Type 2 Diabetes, Inflammatory Bowel Disease and Coronary Heart Disease, all of which have available early intervention or prevention treatments.

## Introduction

In addition to improving our fundamental understanding of basic genetics, phenotypic prediction has obvious practical utility, ranging from crop and livestock applications in agriculture to estimating the genetic component of risk for common human diseases in medicine. For example, a portion of the current guideline on the treatment of blood cholesterol to reduce atherosclerotic cardiovascular risk focuses on estimating a patient’s risk of developing disease^1^; in theory, genetic predictors have the potential to reveal a substantial proportion of this risk early in life (even before clinical risk factors are evident) enabling prophylactic intervention for high-risk individuals. The same logic applies to many other disease areas with available prophylactic interventions including cancers and diabetes.

The field of phenotypic prediction was conceived in plant and animal genetics (reviewed in refs.^2,3^). The first approaches relied on “major genes” – allelic variants of large effect sizes readily detectable by genetic linkage or association. These efforts were quickly followed by strategies adopting polygenic models, most notably the genomic version of the Best Linear Unbiased Predictor (BLUP)^4^.

Similarly, after the early results of human genome-wide association studies (GWAS) became available, the first risk predictors in humans were based on combining the effects of markers significantly and reproducibly associated with the trait, typically those with association statistics exceeding a genome-wide level of significance^5–7^. Almost immediately, after realization that a multitude of small effect alleles play an important role in complex trait genetics^2,3,8^, these methods were extended to accommodate very large (or even all) genetic markers^9–15^. These methods include extensions of BLUP^9,10,16^, or Bayesian approaches that extend both shrinkage techniques and random effect models^11^. Newer methods benefited from allowing for classes of alleles with vastly different effect size distributions. However, these methods require individual level genotype data that do not exist for large meta-analyses and are computationally expensive.

To leverage summary-level data from large-scale GWAS projects, an alternative approach to construct polygenic risk scores based on summary statistics has been introduced^14,17–22,24^. The originally proposed version is additive over genotypes weighted by apparent effect sizes exceeding a given *p*-value threshold. In theory, the risk predictor based on expected true genetic effects given the genetic effects observed in GWAS (conditional mean effects) can achieve the optimal accuracy of linear risk models regardless of underlying genetic architecture by properly down-weighting noise introduced by non-causal variants^23^. In practice, however, implementing the conditional mean predictor poses a dilemma. The GWAS-estimated effect sizes capture genetic effects of all SNPs in linkage disequilibrium (LD), therefore these marginal estimates have to be first deconvoluted into genetic contribution of individual causal SNPs. Furthermore, in order to estimate the conditional mean effects, we need to know the underlying genetic architecture first, but the true architecture is unknown and difficult to model accurately. The current methods circumvent this issue by extensively sampling likely combinations of causal genetic effects under a simplified model of genetic architecture. However, these methods often ignore the correlation of sampling errors of estimated effects between SNPs in LD for the sake of computational efficiency^20,22^. Such approximation can lead to a suboptimal prediction model due to double-counting of correlated sampling errors. In case of dense high-resolution GWAS data, this effect can be severe due to extensive and rank-deficient LD structures. Recent approaches account for correlated sampling errors by applying a Metropolis-Hastings technique to reject proposed states based on a full multivariate likelihood or by assuming a continuous shrinkage prior for the allelic architecture, however their prediction accuracy still depend on the convergence of high-dimensional combinatorial sampling process, and it remains challenging to extend these models to incorporate additional complexity of true architecture^21,24^.

In spite of this methodological complexity, polygenic scores trained on large-scale datasets show some promise for practical applications in medical genetics. Polygenic scores have been used to analyze the UK Biobank, the largest epidemiological cohort that includes genetic data^25^. Individuals with extreme values of polygenic score were shown to have a substantially elevated risk for corresponding diseases, generating enthusiasm for clinical applications of the method.

Here, we propose a novel risk prediction approach called partitioning-based non-parametric shrinkage (NPS). Without specifying a parametric model of underlying genetic architecture, we aim to estimate the conditional mean effects directly from the data. Our method accounts for both types of correlations induced by LD in GWAS summary statistics, namely the correlations of true genetic effects as well as sampling errors, by using eigenvalue decomposition of LD matrix instead of relying on a high-dimensional sampling technique. Despite growing interest in non-parametric prediction models, thus far there has been no non-parametric polygenic score that can fully allow for LD under the conditional mean effect framework^26–31^. We evaluate the performance of this new approach under a simulated genetic architecture of 5 million dense SNPs across the genome. We also test the method using real data in four disease areas: breast cancer, type 2 diabetes, inflammatory bowel disease and coronary heart disease.

## Material and Methods

### Method overview

Our approach is to partition SNPs into groups and determine the relative weights based on predictive value of each partition estimated in the training data (Figure 1A). Intuitively, when there is no LD between SNPs, a partition dominated by non-causal variants will have low power to distinguish cases from controls, whereas the partition enriched with strong signals will be more informative for predicting the phenotype. This is equivalent to approximating the conditional mean effect curve by piecewise linear interpolation. Because of LD, however, we cannot apply the partitioning method directly to GWAS effect sizes. True genetic effects as well as sampling noise are correlated between adjacent SNPs. To prevent estimated genetic signals smearing across partitions, we first transform GWAS data into an orthogonal domain, which we call “eigenlocus” (Figure 1B). Specifically, we use a decorrelating linear transformation obtained by eigenvalue decomposition of the local LD matrix. Both genotypes and sampling errors are uncorrelated in the eigenlocus representation. In this representation, however, true genetic effects do not follow analytically tractable distributions except under infinitesimal and extremely polygenic architectures. Therefore, we apply our partitioning-based non-parametric shrinkage to the estimated effect sizes in the eigenlocus, and then restore them back to the original per-SNP effects.

**Figure 1.**
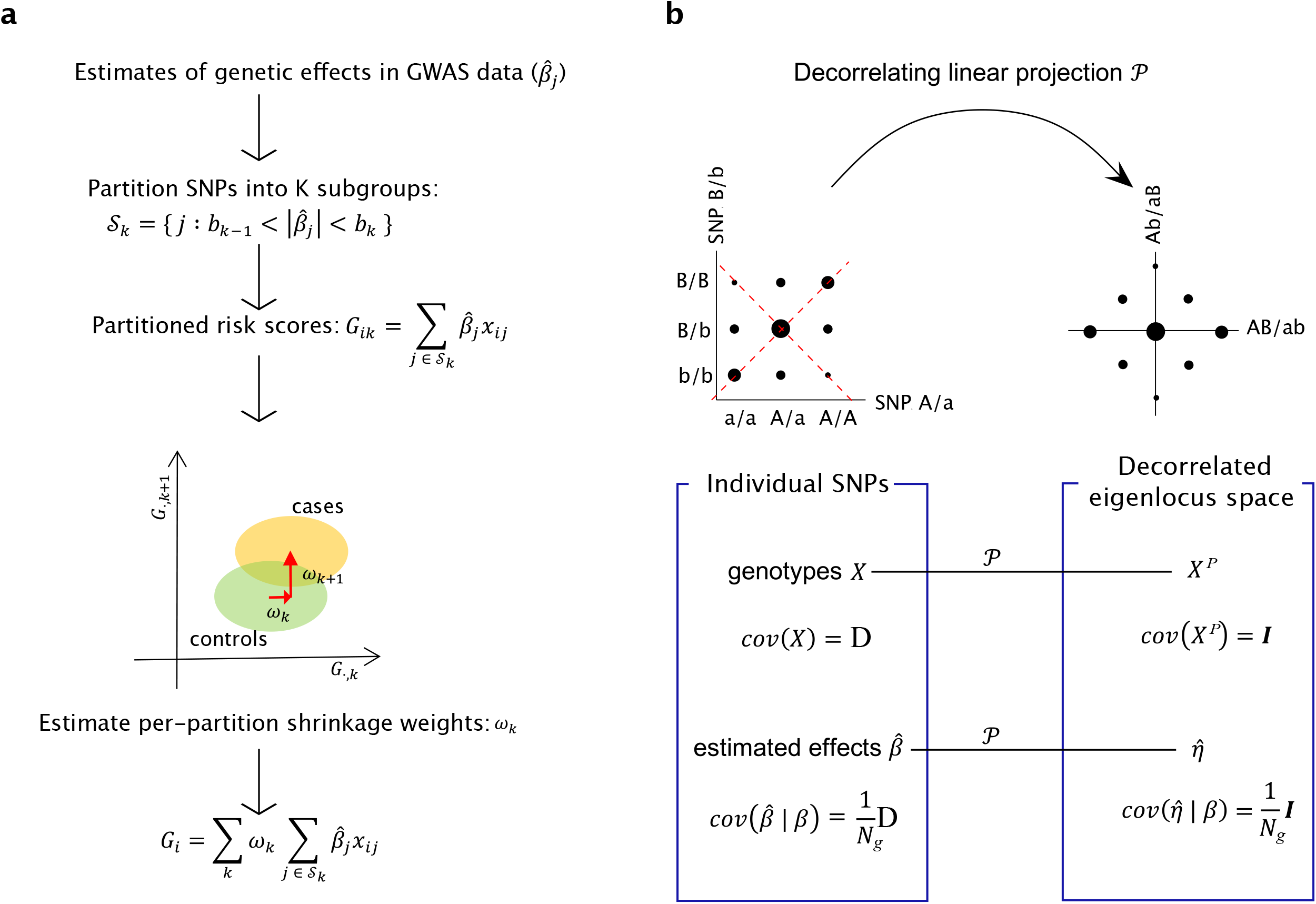
Overview of Non-Parametric Shrinkage (NPS). **(a)** For unlinked markers, NPS partitions SNPs into *K* subgroups splitting the GWAS effect sizes 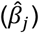 at cut-offs of *b*_0_, *b*_1_, …, *b_K_*. Partitioned risk scores *G_ik_* are calculated for each partition *k* and individual *i* using an independent genotype-level training cohort. The per-partition shrinkage weights *ω_k_* are determined by the separation of *G_ik_* between training cases and controls. Estimating the per-partition shrinkage weights is a far easier problem than estimating per-SNP effects. The training sample size is small but still larger than the number of partitions, whereas for per-SNP effects, the GWAS sample size is considerably smaller than the number of markers in the genome. This procedure “shrinks” the estimated effect sizes not relying on any specific assumption about the distribution of true effect sizes. **(b)** For markers in LD, genotypes and estimated effects are decorrelated first by a linear projection 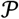 in non-overlapping windows of ~ 2.5 Mb in length, and then NPS is applied to the data. The size of black dots indicates genotype frequencies in population. Before projection, genotypes between SNP 1 and 2 are correlated due to LD (**D**), and thus sampling errors of estimated effects 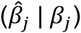 are also correlated between adjacent SNPs. The projection 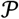 neutralizes both correlation structures. The axes of projection are marked by red dashed lines. *β_j_* denotes the true genetic effect at SNP *j. N_g_* is the sample size of GWAS cohort.

### Decorrelating projection

We split the genome into *L* non-overlapping windows of *m* SNPs each. By default, *m* was set to 4,000 SNPs (~2.5 Mb on average). The window size was chosen to be large enough to capture the majority of LD patterns except near the edge. For the sake of simplicity, we assume that LD is confined to each window and there exists no LD across windows. In each genomic window *l* ∈ {1, …, *L*}, let **X**_*l*_ be an *N* × *m* genotype matrix of *N* individuals and *m* SNPs in the window. We assume that the genotypes are standardized to the mean of 0 and variance of 1. Let 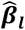 be an *m*-dimensional vector of observed effect sizes from a GWAS and ***β_l_*** be an *m*-dimensional vector of true underlying genetic effects in window *l*. The scales of 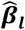 and ***β_l_*** are defined with respect to the standardized genotypes. Then, the LD matrix **D_*l*_** is given by 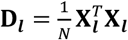 and can be factorized by eigenvalue decomposition into 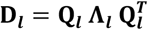, where **Q_*l*_**, is an orthonormal matrix of eigenvectors and **Λ_*l*_**, is a diagonal matrix of eigenvalues.

Now we introduce a linear decorrelating transformation 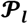, which projects summary statistics 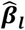 and genotypes **X_*l*_**, into a decorrelated space which we call “**eigenlocus space**.” We call the projection 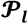 an “**eigenlocus projection**.” 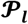 is defined as the following:

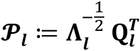

By applying the eigenlocus projection on 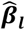, and **X_*l*_**, we obtain the estimated effect sizes 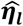 and projected genotypes 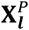 in this eigenlocus space as follows:

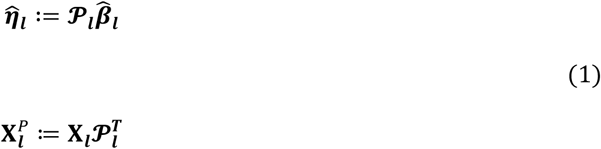

This projection will remove the correlation structure induced by LD in the genotypes 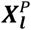 and in the sampling error of estimated effects 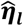. Specifically, in the eigenlocus space, 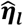 and 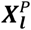 follow the following multivariate normal distributions (See Appendices A and B for the derivation):

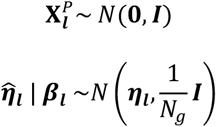

where *N_g_* is the sample size of GWAS from which summary statistics 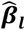 was obtained and ***η_l_*** is the true underlying genetic effect defined by 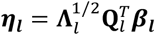.

Due to the rank-deficiency of LD matrix **D_*l*_**, and application of regularization on **D_*l*_**, (described below), the dimension of eigenlocus space *m_l_* can be lower than the total number of SNPs *m* in a given window *l*. Specifically, we set the LD between SNPs to 0 unless the absolute value of estimated LD was greater than 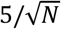. This is to suppress sampling noises in off-diagonal entries of LD matrix. Since the standard error of pairwise LD is approximately 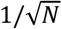 under no correlation, we expect that on average, only 1.7 uncorrelated SNP pairs escape the above regularization threshold in each window. In addition, projections corresponding to eigenvalues less than 0.5 were truncated for the computational efficiency since they were dominated by noises. Although we chose the window size to be large enough to capture the majority of local LD patterns, some LD structures, particularly near the edge, span across windows, which in turn yield cross-window correlations. To eliminate such correlations, we applied LD pruning in the eigenlocus space between adjacent windows. Specifically, we calculated Pearson correlations between projected genotypes belonging to neighboring windows. For the pairs with the absolute Pearson correlation > 0.3, we kept the one yielding a larger absolute effect size and eliminated the other.

By applying the above processing steps in each genomic window *l*, we obtained *m_l_*-dimensional vector of estimated effect sizes 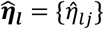 and *N* × *m_l_* matrix of genotypes 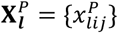 in the eigenlocus space. Here, the index *j* ∈ {1, …, *m_l_*} indicates an individual genetic variation yielded by applying an eigenlocus projection (Eq 1) with eigenvalues **Λ_*l*_** = (*λ_ij_*}. In this representation, we can operate on each genetic variation independently from each other since they are decorrelated.

### Partitioning strategy

Since the SNPs with largest effect sizes span a wide range of values but are sampled only sparsely, we cannot reliably estimate the conditional mean effect for this large-effect tail without assuming a priori parametric assumption on its distribution. This is particularly the case for genome-wide significant SNPs. To solve this problem, we handled the genome-wide significant SNPs as a separate partition from the rest of SNPs and treat them as fixed effect estimates. Specifically, the genome-wide significant SNPs were set aside to a special partition 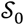, for which the decorrelating projection was set to the identity matrix **I** with eigenvalues of 1. To avoid LD across SNPs in 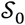, genome-wide significant SNPs were selected into 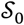 only if the LD between them is low (*r*^2^ < 0.3). Then, we residualized the effects of SNPs in 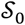 from estimated effects of the rest of SNPs in order to avoid double-counting their genetic effects.

The genetic variants which were not selected to 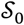 were projected into the eigenlocus space and then grouped into 10 × 10 double-partitions on intervals of eigenvalues *λ_lj_* and absolute estimated effect sizes 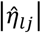. This is because in the eigenlocus space, conditional mean effect 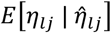 depends not only on the absolute value of estimated genetic effect 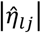 but also on eigenvalue of projection *λ_lj_*. The eigenvalue of projection tracks the scale of true genetic effect in the eigenlocus space (Appendix C). In total, we used 101 partitions in this study including the partition of genome-wide significant SNPs 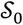.

While fully optimizing the partitioning cut-offs can potentially improve the accuracy of prediction model, this becomes rapidly impractical as the number of partitions increases. NPS requires a large enough number of partitions to closely approximate conditional mean effects, thus the combinatorial search for optimal cut-offs is computationally intractable. Therefore, we applied the following general heuristic, which worked well across our simulation datasets: First, the partitioning cut-offs were selected on the intervals of eigenvalues, equally distributing 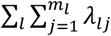 across partitions. This partition scheme evenly distributes the tagged heritability across partitions. The partitions on eigenvalues are denoted here by 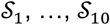 from the lowest to the highest. Then, each partition of eigenvalues 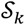 was further partitioned on intervals of 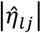, equally distributing 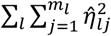 across partitions. This second partitioning scheme is intended to evenly distribute the overall variance in polygenic scores, namely, 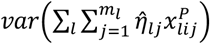, across the partitions. This second partitions of 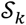 are denoted by 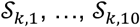 from the lowest to the highest 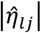.

### Estimation of conditional mean effect

The predicted genetic risk scores of individual *i* ∈ {1, …, *N*} can be represented by the sum of conditional mean effects 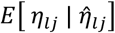 multiplied by genetic dosages 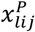 across all genomic windows *l* ∈ {1, …, *L*} and genetic variations *j* ∈ {1, …, *m_l_*} in each window. Instead of deriving conditional mean effects under a genetic architecture prior, we interpolate the conditional mean effects by fitting a linear function 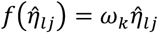 for each partition *k* = 0, …, *K* − 1 as follows:

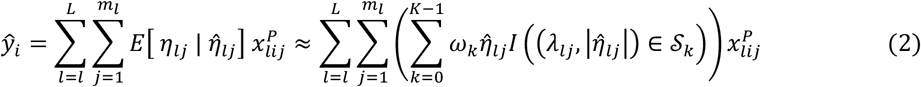

where *I*(·) is an indicator function for the membership of genetic variations to partition *k*, 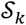 is the set of all genetic variations assigned to partition *k*, and *K* is the total number of partitions, set to 101 by default. The equation (Eq 2) can be further simplified by changing the order of summation as below:

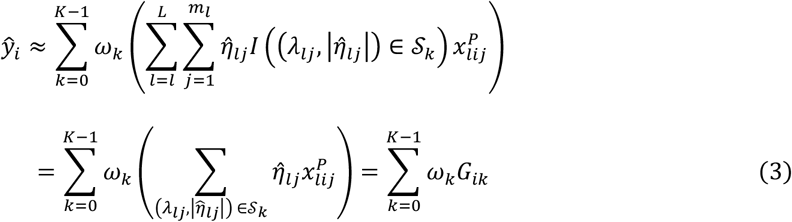

where *G_ik_* is a partitioned polygenic score of individual *i* calculated using only genetic variations belonging to the partition *k*. Then, *ω_k_* becomes equivalent to the per-partition shrinkage weight. Based on (Eq 3), we can estimate *ω_k_* by fitting known phenotypes *y_i_* with partitioned scores *G_ik_* across individuals *i* in a small genotype-level training cohort.

For dichotomous phenotypes without covariates, we used a linear discriminant analysis (LDA) to estimate *ω_k_*. The partitioned scores *G_ik_* calculated in a training cohort form *K*-dimensional feature space, and LDA guarantees the optimal accuracy of the classifier when case and control subgroups follow multivariate normal distributions in the feature space. Since each partition consists of a sufficient number of projected genetic variations, partitioned scores of cases and controls, namely *G_ik_* | *y_i_* follow approximately normal distributions^32^. The variance of partitioned scores is approximately equal between cases and controls since *G_ik_* of an individual partition explains only a small fraction of phenotypic variation on the observed scale in typical GWAS data^33^. Furthermore, due to the decorrelating property of eigenlocus projection, the covariance of *G_ik_* and *G_ik′_* can be assumed to be approximately 0 between different partitions *k* and *k′*. Although in theory, the liability thresholding effect induces slight non-zero covariance between partitions, this effect is typical small and negligible. Thus, LDA-derived shrinkage weights can be independently estimated for each partition and simplify to:

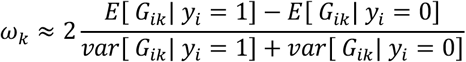

Similarly, for continuous phenotypes or in case of dichotomous traits with covariates, we can estimate per-partition shrinkage weights *ω_k_* by applying the following linear regression model to the training data:

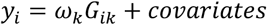

independently for each partition.

In the special case of infinitesimal genetic architecture, in which all SNPs are causal with normally distributed effect sizes, the conditional mean effects has been analytically derived and are predicted to depend only on eigenvalues *λ_lj_*^20^; therefore we can cross-check the accuracy of our shrinkage weights *ω_k_* estimated by NPS in simulations (Appendix D). To apply NPS, we first partitioned genetic variations in the eigenlocus space into 10 subgroups on intervals of their eigenvalues *λ_lj_* as described above but without separating out the genome-wide significant SNPs (Figure 2A). The per-partition shrinkage weights *ω_k_* trained by NPS closely tracked the theoretical optimum in most of the bins. Interestingly, in the lowest and highest partitions of eigenvalues, 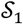 and 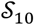, the estimated shrinkage was significantly biased away from the optimal curve. The smallest eigenvalues are too noisy to estimate with the reference LD panel. Therefore, it is correct to down-weight *ω*_1_ almost to 0. In case of partition 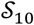, it spans the widest interval of eigenvalues but consists of the fewest number of SNPs. While it is ideal to apply a finer partitioning in this interval so to better interpolate the theoretical curve, the total numbers of SNPs and independent projection vectors in the genome are the fundamental limiting factor.

**Figure 2.**
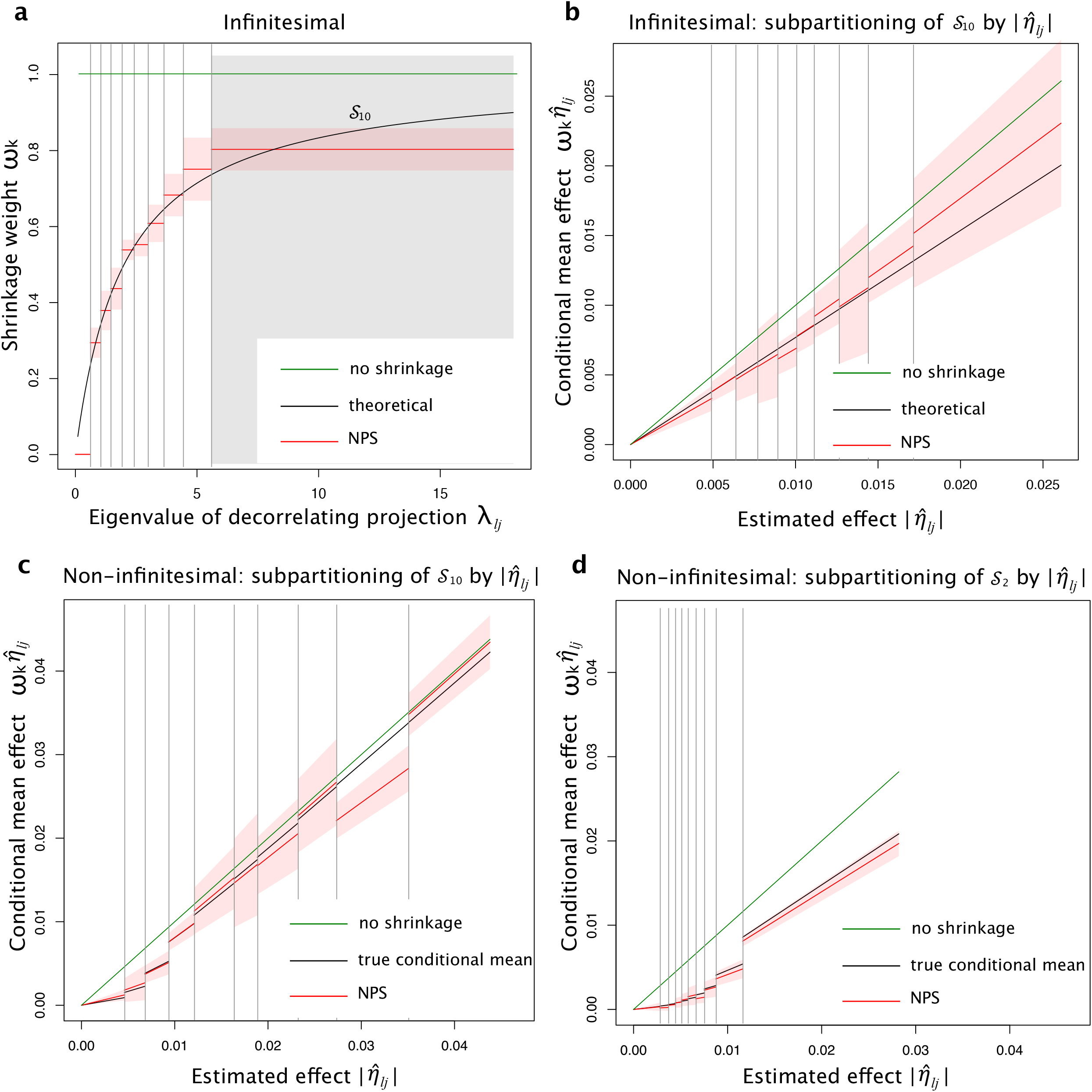
Per-partition shrinkage weights estimated by Non-Parametric Shrinkage (NPS) approximate the conditional mean effects in the decorrelated space. **(a)** NPS shrinkage weights *ω_k_* (red line) compared to the theoretical optimum (black line), 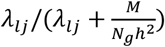, under infinitesimal architecture. The partition of largest eigenvalues 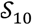 is marked by grey box. **(b)** Conditional mean effects estimated by NPS (red line) in sub-partitions of 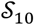 by 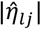, under infinitesimal architecture. The theoretical line (black) is the average over all *λ_lj_*, in 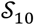. **(c-d)** Conditional mean effects estimated by NPS (red line) in sub-partitions of 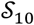 **(c)** and 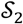 **(d)** on intervals of 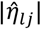 under non-infinitesimal architecture with the causal SNP fraction of 1%. The true conditional means (black) were estimated over 40 simulation runs. **(a-d)** The mean NPS shrinkage weights (red line) and their 95% CIs (red shade) were estimated from 5 replicates. Grey vertical lines indicate partitioning cut-offs. No shrinkage line (green) indicates *ω_k_* = 1. The number of markers *M* is 101,296. The discovery GWAS size *N_g_* equals to *M*. The heritability *h*^2^ is 0.5.

In case of infinitesimal architecture, theory predicts that per-partition shrinkage weights are independent of estimated effect sizes 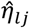. To examine the robustness of NPS, we applied the general 10-by-10 double partitioning on *λ_lj_* and 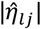 collected under infinitesimal simulations. In overall, the shrinkage weights estimated by double partitioning agree with the theoretical expectation. The estimated conditional mean effects, interpolated with 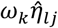, follow the linear trajectory (Figures 2B and S1).

For non-infinitesimal genetic architecture, we do not have an analytic derivation of conditional mean effects; therefore we empirically estimated the conditional means using the true underlying effects *η_lj_* and true LD structure of the population. Here, one percent of SNPs were simulated to be causal with normally distributed effect sizes. As expected, the true conditional mean dips for the lowest values of 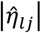 but approaches no shrinkage (*ω_k_* = 1) with increasing values of 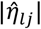 (Figure 2C-D). A notable difference between the partitions of largest eigenvalues and second smallest eigenvalues is that the true conditional mean is very close to no shrinkage for large 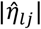 in the former. This is because eigenvalues are proportional to the scale of true effects *η_lj_*; therefore, with large enough eigenvalues, the sampling error becomes relatively small and the estimated effect sizes more accurate. In all partitions, conditional mean effects estimated by NPS stayed very close to the true conditional means (Figure S2).

### Back-conversion from the eigenlocus space to per-SNP effects

Rewriting the equation (Eq 2) using matrix operations, we can reformulate the *N*-dimensional vector of predicted genetic risk scores 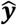 using the original SNP genotypes **X_*l*_** instead of eigenlocus genotypes 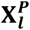 as follows:

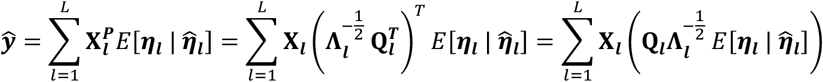

from the definition of 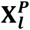 (Eq 1). We obtain the conditional mean effects by non-parametric shrinkage in the following form:

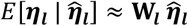

where **W_*l*_**, is an *m_l_* × *m_l_* diagonal matrix with diagonal entries {*w_jj_*} defined as:

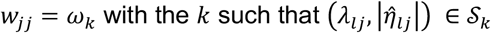

where *k* is the partition to which the *j*-th projected genetic variation belong in the eigenlocus space. Therefore, the reweighted effects in the original per-SNP scale can be retrieved back by computing 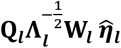.

### Application of NPS to genome-wide datasets

The estimated effect size at each SNP is available as summary statistics from a large discovery GWAS study. As these estimated effects were represented as per-allele effects, we converted them relative to standardized genotypes by multiplying by 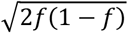, where *f* is the allele frequency of each SNP in the discovery GWAS cohort.

Because the accuracy of eigenlocus projection declines near the edge of windows, the overall performance of NPS is affected by the placement of window boundaries relative to locations of strong association peaks. To alleviate such dependency, we repeated the same NPS procedure shifting by 1,000, 2,000, and 3,000 SNPs and took the average reweighted effect sizes across four NPS runs. When NPS was run in parallel on up to 88 processors (22 chromosomes x 4 window shifts), it took total computation time of 3 to 6 hours for each dataset.

### Simulation of genetic architecture with dense genome-wide markers

For simulated benchmarks, we generated genetic architecture with 5 million dense genome-wide markers from the 1000 Genomes Project. We kept only SNPs with MAF > 5% and Hardy-Weinberg equilibrium test *p*-value > 0.001. We used EUR panel (n=404) to populate LD structures in simulated genetic data. Due to the limited sample size of the LD panel, we regularized the LD matrix by applying Schur product with a tapered banding matrix so that the LD smoothly tapered off to 0 starting from 150 kb up to 300 kb^34^.

Next, we generated genotypes across the entire genome, simulating the genome-wide patterns of LD. We assume that the standardized genotypes follow a multivariate normal distribution. Since we assume that LD travels no farther than 300 kb, as long as we simulate genotypes in blocks of length greater than 300 kb, we can simulate the entire chromosome without losing any LD patterns by utilizing a conditional multivariate normal distribution as the following. The genotypes for the first block of 1,250 SNPs (average 750 kb in length) were sampled directly out of multivariate normal distribution *N*(*μ* = 0, Σ = **D**_1_). From the next block, we sampled the genotypes of 1,250 SNPs each, conditional on the genotypes of previous 1,250 SNPs. When the genotype of block *l* is x_*l*_ and the LD matrix spanning block *l* and *l* + 1 is split into submatrices as the following:

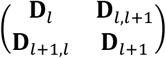

then, the genotype of next block *l* + 1 follows a conditional MVN as:

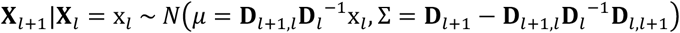

After the genotype of entire chromosome was generated in this way, the standardized genotype values were converted to allelic genotypes by taking the highest *nf* and lowest *n*(1 − *f*)^2^ genotypes as homozygotes and the rest as heterozygotes under the Hardy-Weinberg equilibrium. *n* is the number of simulated samples, and *f* is the allele frequency of each SNP. This MVN-based simulator can efficiently generate a very large cohort with realistic LD structure across the genome and guarantees to produce homogenous population without stratification.

We simulated three different sets of genetic architecture: point-normal mixture, MAF dependency and DNase I hypersensitive sites (DHS). The point-normal mixture is a spike-and-slab architecture in which a fraction of SNPs have normally distributed causal effects *β_j_* for SNP *j* as below:

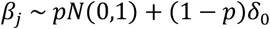

where *p* is the fraction of causal SNPs being 1, 0.1 or 0.01% and *δ*_0_ is a point mass at the effect size of 0. For the MAF-dependent model, we allowed the scale of causal effect sizes to vary across SNPs in proportion to 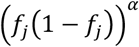 with *α* = −0.25^35^ as follows:

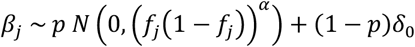

Finally, for the DHS model, we further extended the MAF-dependent point-normal architecture to exhibit clumping of causal SNPs within DHS peaks. Fifteen percent of simulated SNPs were located in the master DHS sites that we downloaded from the ENCODE project. We assumed a five-fold higher causal fraction in DHS (*p_DHS_*) compared to the rest of genome in order to simulate the enrichment of per-SNP heritability in DHS reported in the previous study^36^. Specifically, *β_j_* was sampled from the following distribution:

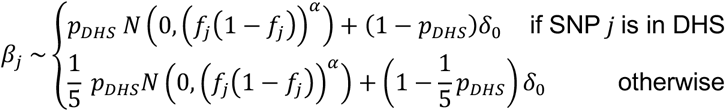

In each genetic architecture, we simulated phenotypes for discovery, training and validation populations of 100,000, 50,000 and 50,000 samples, respectively, using a liability threshold model of the heritability of 0.5 and prevalence of 0.05. In the discovery population, we obtained GWAS summary statistics with Plink by testing for the association with the total liability instead of case/control status; this is computationally easier than to generate a large case/control GWAS cohort directly, and the estimated effect sizes are approximately equivalent by a common scaling factor. With the prevalence of 0.05, statistical power of quantitative trait association studies using the total liability is roughly similar to those of dichotomized case/control GWAS studies of same sample sizes^37^. For the training dataset, we assembled a cohort of 2,500 cases and 2,500 controls by down-sampling controls out of the simulated population of 50,000 samples. The validation population was used to evaluate the accuracy of prediction model in terms of *R*^2^ of the liability explained and Nagelkerke’s *R*^2^ to explain case/control outcomes.

### GWAS summary statistics

GWAS summary statistics are publicly available for phenotypes of breast cancer^38,39^, inflammatory bowel disease (IBD)^40^, type 2 diabetes (T2D)^41^ and coronary artery disease (CAD)^42^. These GWAS summary statistics were based only on Caucasian samples with an exception of CAD, for which 13% of discovery cohort comprised of non-European ancestry.

### UK Biobank

UK Biobank samples were used for training and validation purposes. Case and control samples were defined as follows. Breast cancer cases were identified by ICD10 codes of diagnosis. Controls were selected from females who were not diagnosed with or did not self-report history of breast cancer. We excluded individuals with history of any other cancers, *in situ* neoplasm or neoplasm of unknown nature or behavior from both cases and controls. For IBD, we identified case individuals by ICD10 or self-reported disease codes of Crohn’s disease, ulcerative colitis or IBD. Controls were randomly selected excluding participants with history of any auto-immune disorders. For T2D, cases were identified by ICD10 diagnosis codes or by questionnaire on history of diabetes combined with the age of diagnosis over 30. However, our T2D cases may include a small fraction of Type 1 Diabetic cases misdiagnosed as T2D (3.7%) as previously reported^43^. For early-onset CAD, case individuals were identified by ICD10 codes of diagnosis or cause of death. The early-onset was determined by the age of heart attack on the questionnaire (≤ 55 for men and ≤ 65 for women). Individuals with history of CAD were excluded from controls regardless of the age of onset. The latest CAD summary statistics include UK Biobank samples in the interim release; thus, to avoid sample overlap, we used only post-interim samples, which were identified by genotyping batch IDs. For all phenotypes, our case definition includes both prevalent and incident cases.

For genotype QC, we filtered out SNPs with MAF below 5% or INFO score less than 0.4. We also excluded tri-allelic SNPs and InDels. For all phenotypes, we filtered out participants who were retracted, not from white British ancestry, or had indication of any QC issue in UK Biobank. We included only samples which were genotyped with Axiom array. Related samples were excluded to avoid potential confounding. The samples were randomly split to training and validation cohorts. Controls were down-sampled to the case to control ratio of 1:1 to assemble training cohorts, but no down-sampling was applied to validation cohorts to keep the original case prevalence.

### Partners Biobank

We used Partners Biobank^44^ to evaluate the accuracy of prediction models in an independent validation cohort. These genotyping data were previously generated using the MEGA-Ex array. Markers with monomorphic allele frequency, complementary alleles, less than 99.5% genotyping rate, or deviation from Hardy-Weinberg equilibrium (*P* < 0.05) were removed. Then, statistical imputation was conducted to infer genotypes at missing markers using Eagle v2.4 and IMPUTE v4 on the reference panel (1000 Genomes Phase 3). Excluding samples of non-European ancestry, a total of 16,839 samples from US white population were available for use. Participants with breast cancer, IBD, T2D and CAD were identified using a phenotype query algorithm with the PPV parameter of 0.90^45^. To obtain early-onset CAD, both cases and controls were restricted to men with age ≤ 55 and women with age ≤ 65. Since the prevalence of early-onset CAD and T2D are sex-dependent, we included the sex covariate in the genetic risk model for CAD and T2D. For all methods, the coefficient of sex covariate was estimated in the training cohort of UK Biobank.

### LDPred

The accuracy of LDPred was evaluated in simulated and real datasets using the default parameter setting. The underlying causal fraction parameter was optimized using the training cohort, which is available as individual-level genotype data. Specifically, the causal SNP fractions of 1, 0.3, 0.1, 0.03, 0.01, 0.003, 0.001, 0.0003 and 0.0001 were tested in the training data, and the prediction model yielding the highest prediction *R*^2^ was selected for validation. The training genotypes were also used as a reference LD panel.

LDPred accepts only hard genotype calls as inputs at the training step. Thus, for real data we converted imputed allelic dosages to most likely genotypes after filtering out SNPs with genotype probability < 0.9. SNPs with the missing rate > 1% or deviation from Hardy-Weinberg equilibrium (*P* < 10^−5^) were also excluded. Prediction models were trained using only SNPs which passed all QC filters in both training and validation datasets, as recommended by the authors. SNPs with complementary alleles were excluded automatically by LDPred. In simulations, all genotypes were generated as hard calls, and complementary alleles were avoided; thus, the exactly same set of SNPs were used for both LDPred and NPS. In a subset of datasets, we further examined the accuracy of LDPred when it was run only with directly genotyped SNPs. In simulated datasets, we assumed that both training and validation cohorts were genotyped with Illumina HumanHap550v3 array, restricting the genotype data to 490,504 common SNPs. For UK Biobank datasets, prediction models were constrained to up to 354,110 common SNPs in UK Biobank Axiom array. In the case of validation in Partners Biobank, we did not consider running LDPred only with genotyped SNPs since too few SNPs were directly genotyped in both UK Biobank and Partners Biobank; thus, we validated LDPred only using overlapping markers in imputed data of two cohorts.

### LD Pruning and Thresholding

LD Pruning and Thresholding (P+T) algorithm was evaluated using PRSice software in the default setting^46^. In real data, imputed allelic dosages were converted to hard-called genotypes similarly as for LDPred. A training cohort was used as a reference LD panel and to optimize pruning and thresholding parameters. The best prediction model suggested by PRSice was evaluated in validation cohorts.

### PRS-CS

PRS-CS algorithm was benchmarked using the default parameter setting^24^. The optimal *ϕ* parameter values were optimized in training cohorts, and the highest performing model was evaluated in validation cohorts. For the reference LD panel, we used a set of simulated genotypes produced by our MVN simulator in order to accurately capture the underlying LD structure of our simulated datasets; in real data, we used the “EUR” reference LD panel provided in the software. Imputed allelic dosages were converted to hard-called genotypes similarly as recommended by the authors.

## Results

### Application to simulated data

To benchmark the accuracy of NPS, we simulated the genetic architecture using the real LD structure of 5 million dense common SNPs from the 1000 Genomes Project (Methods). We considered the causal fraction of SNPs from 1% to 0.01%, dependency of heritability on minor allele frequency (MAF) and enrichment of heritability in DNase I hypersensitive sites (DHS) based on the previous literature^35,36,47^. The prediction accuracy of NPS remained robust across the simulated genetic architectures (Table 1 and Table S1). We measured prediction accuracy using Nagelkerke ***R*^2^** and odds ratio at the highest 5% tail of the polygenic score distribution. The latter measure has been popularized by a recent study that reported that the tails of the polygenic score distribution are associated with risk that is similar to monogenic mutations^25^.

**Table 1.** Comparison of prediction accuracy in simulated genetic architecture.

| % causal SNPs | Method | $R^2_{\text{Nagelkerke}}$ | % $h^2$ explained | 5% Tail OR | NPS $R^2_{\text{Nag}}$ compared to | | |
| --- | --- | --- | --- | --- | --- | --- | --- |
|  |  |  |  |  | P+T | LDPred | PRS-CS |
| 1% | P+T | 0.050 | 14.8 | 3.18 |  |  |  |
|  | LDPred | 0.068 | 20.6 | 3.66 |  |  |  |
|  | PRS-CS | 0.075 | 22.0 | 4.02 |  |  |  |
|  | NPS | 0.085 | 24.6 | 4.27 | 1.68 * | 1.25 * | 1.13 * |
| 0.1% | P+T | 0.136 | 40.8 | 6.32 |  |  |  |
|  | LDPred | 0.080 | 23.0 | 4.08 |  |  |  |
|  | PRS-CS | 0.156 | 44.8 | 7.03 |  |  |  |
|  | NPS | 0.179 | 51.2 | 8.09 | 1.31 * | 2.22 * | 1.14 * |
| 0.01% | P+T | 0.213 | 61.4 | 9.92 |  |  |  |
|  | LDPred | 0.153 (0.268)<sup>a</sup> | 43.8 (74.6)<sup>a</sup> | 7.66 (13.37)<sup>a</sup> |  |  |  |
|  | PRS-CS | 0.228 | 65.3 | 10.35 |  |  |  |
|  | NPS | 0.328 | 92.6 | 17.19 | 1.54 * | 2.14 * | 1.44 * |

Non-parametric shrinkage (NPS) is more robust and accurate compared to other methods in simulated datasets. The simulations incorporate the dependency of heritability on minor allele frequency and clumping of causal SNPs in known DHS elements. The heritability was 0.5, and the prevalence was 5%. The number of markers was 5,012,500. The GWAS sample size was 100,000. Prediction models were optimized in the training cohort of 2,500 cases and 2,500 controls. $R^2$ of prediction was measured in the validation cohort of 50,000 samples. The $h^2$ explained stands for the proportion of heritability on the liability scale explained by polygenic scores. The star (\*) indicates a significant improvement in Nagelkerke's $R^2$ (paired t-test; $P < 0.05$ ). <sup>a</sup>The accuracy of LDPred varies widely depending on the convergence of prediction model; thus, we report the maximum $R^2$ in parenthesis as well as the average performance.

We evaluated the performance of NPS vis-a-vis two popular methods LDPred and P+T, as well as the newest method PRS-CS with the superior reported accuracy^13,20,24^ (Tables S1–5). LDPred is the state-of-the-art Bayesian parametric method, which is similarly based on summary statistics estimated in large GWAS datasets and an independent training set with individual-level data. PRS-CS is a new sophisticated extension of the Bayesian strategy. We found that our method resulted in more accurate predictions than all three methods across a range of genome-wide simulations. PRS-CS was shown to be more accurate than P+T and LDPred on simulated data, although less accurate than NPS. The improvement over LDPred is seemingly surprising given that some of the simulated allelic architectures are the spike-and-slab allelic architecture for which LDPred is expected to be optimal as a Bayesian method. However, we found that in most simulations, LDPred adopted the infinitesimal or extremely polygenic model irrespective of the true simulated regime, pointing to the challenge of computational optimization in the parametric case (Table S3). The simulations suggest that the well-optimized parametric models are capable of generating good predictions, but NPS is much more robust and does not suffer from optimization issues. Overall, NPS improves accuracy consistently for all simulated allelic architectures for both Negelkerke ***R*^2^** and odds ratios at 5% tail (Table 1).

### Application to real data

We benchmarked the accuracy of NPS and other methods using publicly available GWAS summary statistics and training and validation cohorts assembled with UK Biobank samples (Methods)^38–42,48^. For all three phenotypes except coronary artery disease, NPS showed significantly higher accuracy than LDPred or P+T (Table 2, Tables S6–9 and Figures S3–7) and highly similar (statistically indistinguishable) accuracy compared to PRS-CS. In particular, our method and PRS-CS outperformed the other two methods by greater magnitudes with more recent GWAS summary statistics with finer resolution. For example, the latest breast cancer GWAS study has twice as large sample size as the previous study and used a custom genotyping array to densely genotype known cancer susceptibility loci. The *R*^2^ of our method increased by 1.5-fold with the latest breast cancer data whereas the accuracy of LDPred did not improve at all. The *R*^2^ of P+T increased by 1.25-fold, but the gain is mainly due to the inferior accuracy with older GWAS data.

**Table 2.** Accuracy of polygenic prediction in real data.

| Discovery GWAS | Training<br>(UK Biobank) | Validation<br>(UK Biobank) | Method | $R^2_{Nag}$ | 5% Tail OR |
| --- | --- | --- | --- | --- | --- |
| Breast Cancer 2015<br>(N=~120,000) | N=3,956/3,956 | N=3,957/73,652 | P+T | 0.021 | 2.28 |
|  |  |  | LDPred | 0.026 | 2.42 |
|  |  |  | PRS-CS | 0.030 | 2.60 |
|  |  |  | NPS | 0.030 | 2.53 |
| Breast Cancer 2017<br>(N=~230,000) |  |  | P+T | 0.027 | 2.37 |
|  |  |  | LDPred | 0.026 | 2.33 |
|  |  |  | PRS-CS | 0.043 | 2.96 |
|  |  |  | NPS | 0.045 | 3.01 |
| Inflammatory Bowel<br>Disease<br>(N=~35,000) | N=2,483/2,483 | N=2,482/157,272 | P+T | 0.028 | 3.00 |
|  |  |  | LDPred | 0.027 | 2.77 |
|  |  |  | PRS-CS | 0.040 | 3.67 |
|  |  |  | NPS | 0.035 | 3.60 |
| Type 2 Diabetes<br>(N=~160,000) | N=7,298/7,298 | N=7,298/144,020 | P+T | 0.046 | 3.04 |
|  |  |  | LDPred | 0.059 | 3.51 |
|  |  |  | PRS-CS | 0.066 | 3.99 |
|  |  |  | NPS | 0.065 | 3.81 |
| Coronary Artery<br>Disease<br>(N=~330,000) | N=2,000/2,000 | N=773/62,512 | P+T | 0.063 | 5.17 |
|  |  |  | LDPred | 0.078 | 5.65 |
|  |  |  | PRS-CS | 0.075 | 4.92 |
|  |  |  | NPS | 0.073 | 5.21 |

Non-Parametric Shrinkage (NPS) and PRS-CS outperform both Pruning and Thresholding (P+T) and LDPred in real data. Both training and validation cohorts were sampled from UK Biobank. The tail Odds Ratio (OR) stands for the odds ratios of cases over controls at the 5% tail in polygenic score distribution compared to the rest. For CAD and T2D, all prediction models were trained and validated with the sex covariate to account for the difference of disease prevalence by sex.

Since our method estimates a large number of parameters from the training data, it might be particularly vulnerable to overfitting cryptic genetic features common to both training and testing data which may result in inflated prediction accuracy. To eliminate this possibility, we benchmarked the prediction models in Partners Biobank, as an independent validation cohort (Methods)^44^. For all phenotypes, NPS outperformed both P+T and LDPred and showed similar accuracy as PRS-CS (Table 3 and Tables S10–13). NPS also has a higher odds ratio at 5% distribution tail than PRS-CS consistently for all phenotypes, although this improvement is not statistically significant (Table 3).

**Table 3.** Accuracy of polygenic prediction in independent validation cohorts.

| Discovery GWAS | Training<br>(UK Biobank) | Validation<br>(Partners) | Method | $R^2_{Nag}$ | 5% Tail OR |
| --- | --- | --- | --- | --- | --- |
| Breast Cancer 2017<br>(N~230,000) | N=3,956/3,956 | N=754/8,324 | P+T | 0.016 | 1.56 |
|  |  |  | LDPred | 0.015 | 1.78 |
|  |  |  | PRS-CS | 0.034 | 2.23 |
|  |  |  | NPS | 0.034 | 2.32 |
| Inflammatory<br>Bowel Disease<br>(N~35,000) | N=2,483/2,483 | N=839/16,000 | P+T | 0.050 | 3.57 |
|  |  |  | LDPred | 0.038 | 3.07 |
|  |  |  | PRS-CS | 0.065 | 4.11 |
|  |  |  | NPS | 0.069 | 4.32 |
| Type 2 Diabetes<br>(N~160,000) | N=7,298/7,298 | N=2,026/14,813 | P+T | 0.038 | 2.10 |
|  |  |  | LDPred | 0.046 | 2.51 |
|  |  |  | PRS-CS | 0.058 | 2.80 |
|  |  |  | NPS | 0.054 | 2.97 |
| Coronary Artery<br>Disease<br>(N~330,000) | N=2,000/2,000 | N=268/7,107 | P+T | 0.018 | 2.72 |
|  |  |  | LDPred | 0.016 | 2.31 |
|  |  |  | PRS-CS | 0.027 | 3.16 |
|  |  |  | NPS | 0.025 | 4.10 |

## Discussion

Understanding how phenotype maps to genotype has always been a central question of basic genetics. With the explosive growth in the amount of training data, there is also a clear prospect and enthusiasm for clinical applications of the polygenic risk prediction^25,49^. The current reality is, however, that most large-scale GWAS datasets are available in the form of summary statistics only. Nonetheless, data on a limited number of cases are frequently available from epidemiological cohorts such as UK Biobank or from public repositories with a secured access such as dbGaP. This motivated us to develop a method that is primarily based on summary statistics but also benefits from smaller training data at the raw genotype resolution. Although we heavily rely on the training data to construct a prediction model, the requirement for out-of-sample training data is not unique for our method. Widely-used thresholding-based polygenic scores and Bayesian parametric methods also need genotype-level data to optimize their model parameters^20,50^. Also, our method assumes – similar to other methods – that all datasets come from a homogeneous population. It has been shown that polygenic risk models are not transferrable between populations due to differences in allele frequencies and patterns of linkage disequilibrium^51^, which is a problem that should be addressed by future work in this field.

Human phenotypes vary in the degree of polygenicity^52^, in the fraction of heritability attributable to low-frequency variants^35^ and in other aspects of allelic architecture^47,53^. The optimality of a Bayesian risk predictor is not guaranteed when the true underlying genetic architecture deviates from the assumed prior. In particular, recent studies have revealed complex dependencies of heritability on minor allele frequency (MAF) and local genomic features such as regulatory landscape and intensity of background selections^35,36,47,52,53^. Several studies have proposed to extend polygenic scores by incorporating additional complexity into the parametric Bayesian models, yet these methods were not applied to genome-wide sets of markers due to computational challenges^54,55^. Recently, there has been a growing interest in non-parametric or semi-parametric approaches, such as those based on modeling of latent variables or kernel-based estimation of prior or marginal distributions, however, thus far they cannot leverage summary statistics or directly account for the linkage disequilibrium structure in the data^26–29^. To address these issues, we developed NPS, a non-parametric method which is agnostic to allelic architecture. In simulations, we show that this approach should be advantageous across a wide range of phenotypes and traits with differing underlying architectures and find that it outperforms existing prediction methods in UK Biobank for four different traits of medical interest. NPS is flexible to incorporate additional complexity of true genetic architecture. Our non-parametric approach has been recently adopted by LDPred-funct, an extension of LDPred to incorporate functional annotations^56^. Finally, as demonstrated in the prediction accuracy using two different breast cancer GWAS summary statistics, with increasing size and marker density in case-control association studies across a range of diseases, our NPS method should outperform traditional parametric approaches for identifying individuals at increased risk.

## Appendix

### A. Distribution of projected genotypes in the eigenlocus space

Let *X_i_* be an *m*-dimensional genotype vector of all SNPS in genomic window *l* and individual *i*. We drop the subscript for genomic window for the sake of simplicity when it is clear from the context. The standardized genotype *X_i_* follows the following multivariate normal distribution:

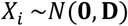

where **D** is a LD matrix of the window. Since the projected genotype 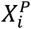 is derived by applying eigenlocus projection 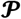 on *X_i_* by definition (Eq 1), 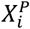 also follows a multivariate normal distribution. Specifically, the distribution of 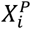 is:

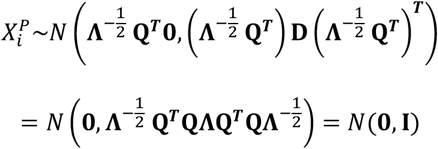

since **D = QΛQ^*T*^** and **Q^*T*^ Q = I**. The projected genotypes in the eigenlocus space are decorrelated with the covariance of **I**.

### B. Distribution of effect size estimates in the eigenlocus space

In the discovery GWAS, the estimated effect sizes 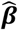 are calculated by linear regression as below:

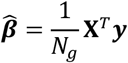

where ***y*** is an *N_g_*-dimensional phenotype vector and *N_g_* is the sample size of GWAS cohort. For convenience, we assume that ***y*** is standardized to the mean of 0 and variance of 1. At this time, we treat genotypes as fixed variables and model the true underlying genetic effects ***β*** and residuals ***ϵ*** as random. Since ***y* = X*β* + *ϵ***,

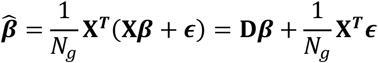

where the residual ***ϵ*** follows an *N_g_*-dimensional multivariate normal distribution 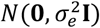. In an individual window, the genetic effects explain only a small fraction of phenotypic variation, therefore we can assume that 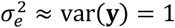. The distribution of sampling noise in 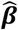, namely the distribution of 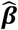 given ***β***, follows:

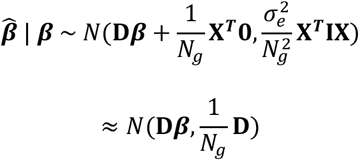

since 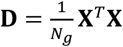. Since the estimated effect size 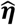 in the eigenlocus space is obtained by applying 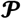 on 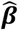 by definition (Eq 1), the distribution of 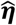 given ***β*** also follows a multivariate normal distribution:

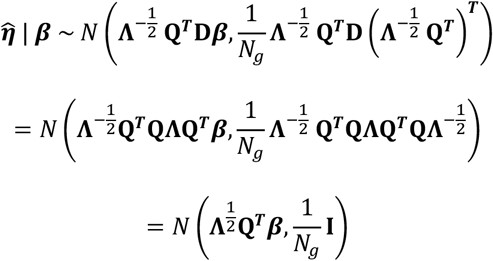

since **D = QΛQ^*T*^** and **Q^*T*^ Q = I**. The sampling noise in 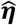 is now decorrelated with the covariance of 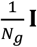. Hence, the eigenlocus projection 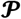 removes correlations in both genotypes and sampling noise of effect size estimates.

### C. Interpretation of eigenvalues

Let ***β*** be the *m*-dimensional vector of true genetic effect at *m* SNPs in a genomic window. We assume that ***β*** is symmetric at 0 and independent at each SNP. Then, the distribution of true genetic effects ***η*** = {*η_j_*} in the eigenlocus space will follow:

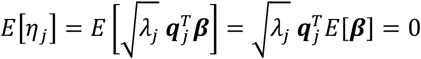

where *λ_j_* and ***q**_j_* are the eigenvalue and eigenvector, respectively, projecting ***β*** to *η_j_* by (Eq 1). If we put that eigenvector ***q**_j_* is (*q*_1*j*_ … *q_mj_*)^*T*^ and ***β*** is (*β*_1_ … *β_m_*)^*T*^, the variance of true genetic effects for an eigenlocus is:

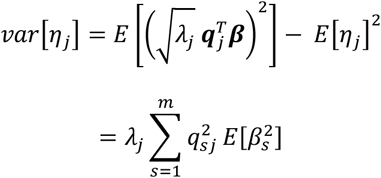

Therefore, in general, *var*[*η_j_*], is directly proportional to eigenvalue *λ_j_*. In particular, when all SNPs have the same variance of per-SNP effect sizes 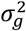,

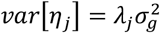

since 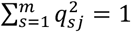.

### D. Conditional mean effects under infinitesimal genetic architecture in the eigenlocus space

Under infinitesimal genetic architecture, the conditional mean effect has been analytically derived by Vilhjalmsson et al.^20^:

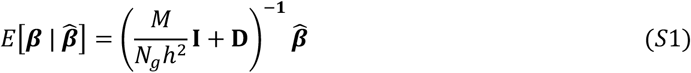

where *N_g_* is the sample size of GWAS cohort, *h*^2^ is the heritability of trait, *M* is the total number of SNPs, and **D** is the LD matrix of full rank. Then, **D** can be factorized into **D = QΛQ^*T*^** with eigenvalues **Λ** and eigenvectors **Q**. Since

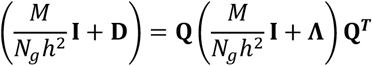

and

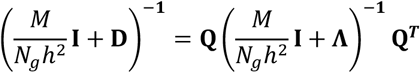

we can reformulate the equation (Eq S1) as follows:

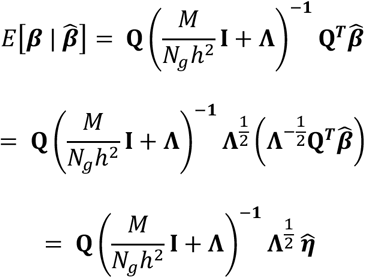

by the definition of 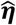 (Eq 1). Hence,

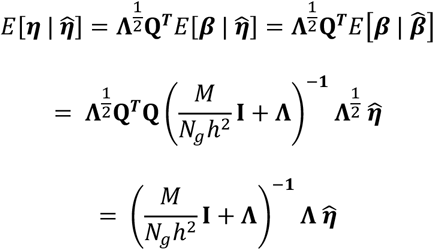

by the definition of ***η***. Therefore, for the *j*-th eigenlocus projection defined by eigenvalue *λ_j_* and eigenvector ***q_j_***, the conditional mean effect is given as the following:

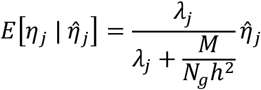

Thus, under infinitesimal architecture, the conditional mean effect 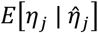 simplifies to 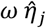, where *ω* is the theoretically optimal shrinkage weight and depends only on eigenvalues as follow:

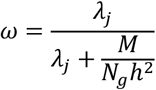

## Supplemental Data

Supplemental Data include seven figures and thirteen tables.

## Acknowledgements

NOS was supported in part by grants K08HL114642, R01HL131961, UM1HG008853, and by The Foundation for Barnes-Jewish Hospital. SK was supported by a Research Scholar award from the Massachusetts General Hospital, the Donovan Family Foundation, R01HL107816, a grant from Fondation Leducq, and an investigator-initiated grant from Merck. SRS was supported by R35GM127131, R01MH101244 and U01HG006500. This research has been conducted using the UK Biobank Resource under Application Number 31063.

## Web Resources

The NPS software is available at http://github.com/sgchun/nps/.

---

**Figure S1.**
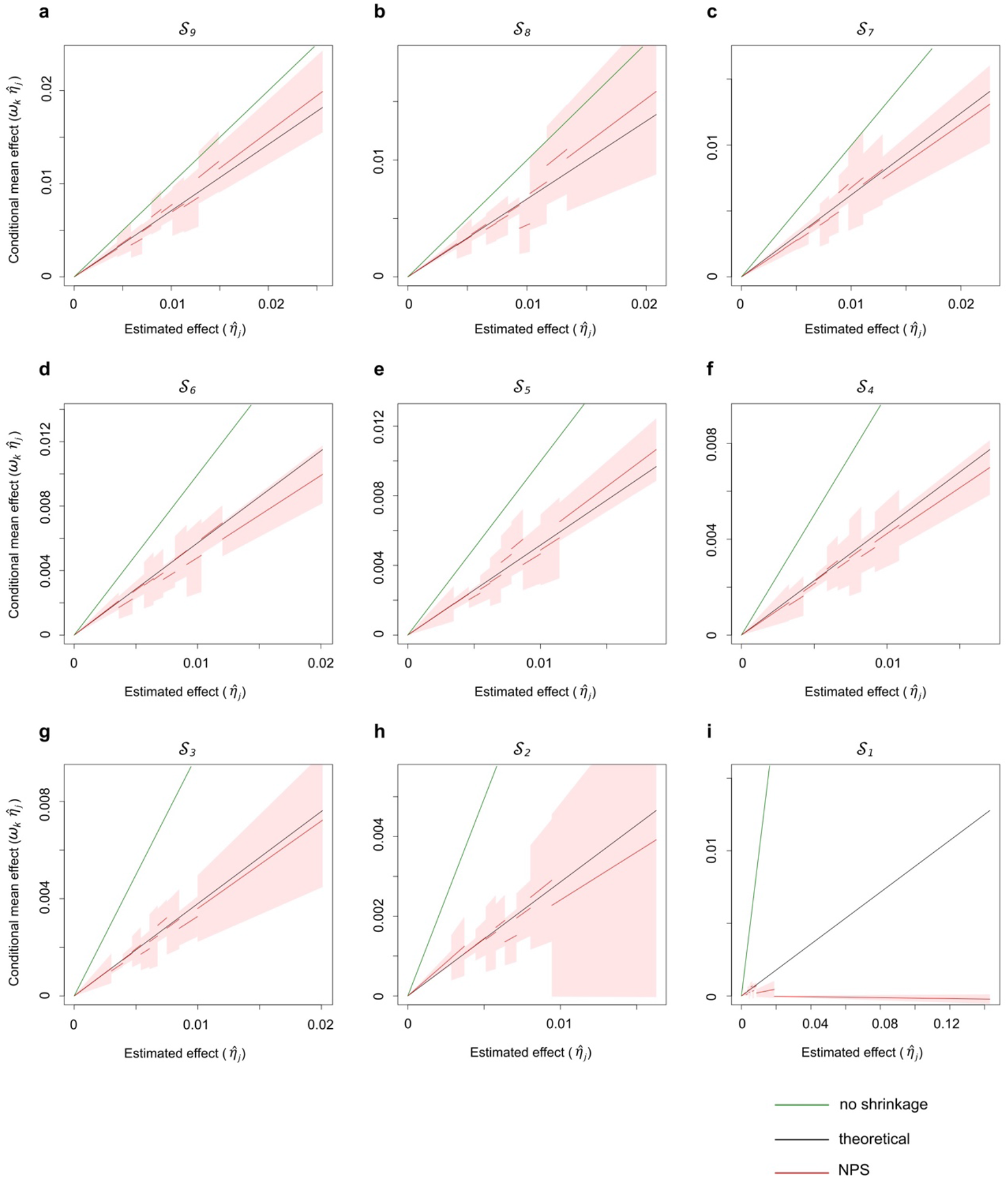
NPS approximates the conditional mean effects: infinitesimal genetic architecture 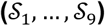. NPS shrinkage weights *ω_k_* (red line) are compared to the theoretical optimum (black line), 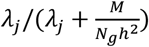, under the infinitesimal architecture. 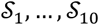 indicate the partitions of lowest to highest eigenvalues of projection. The mean NPS shrinkage weights (red line) and their 95% CIs (red shade) were estimated from 5 replicates. No shrinkage line (green) indicates *ω_k_* = 1. The number of markers *M* is 101,296. The discovery GWAS size *N* equals to *M*. The heritability *h*^2^ is 0.5. See Figure 2B for 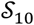.

**Figure S2.**
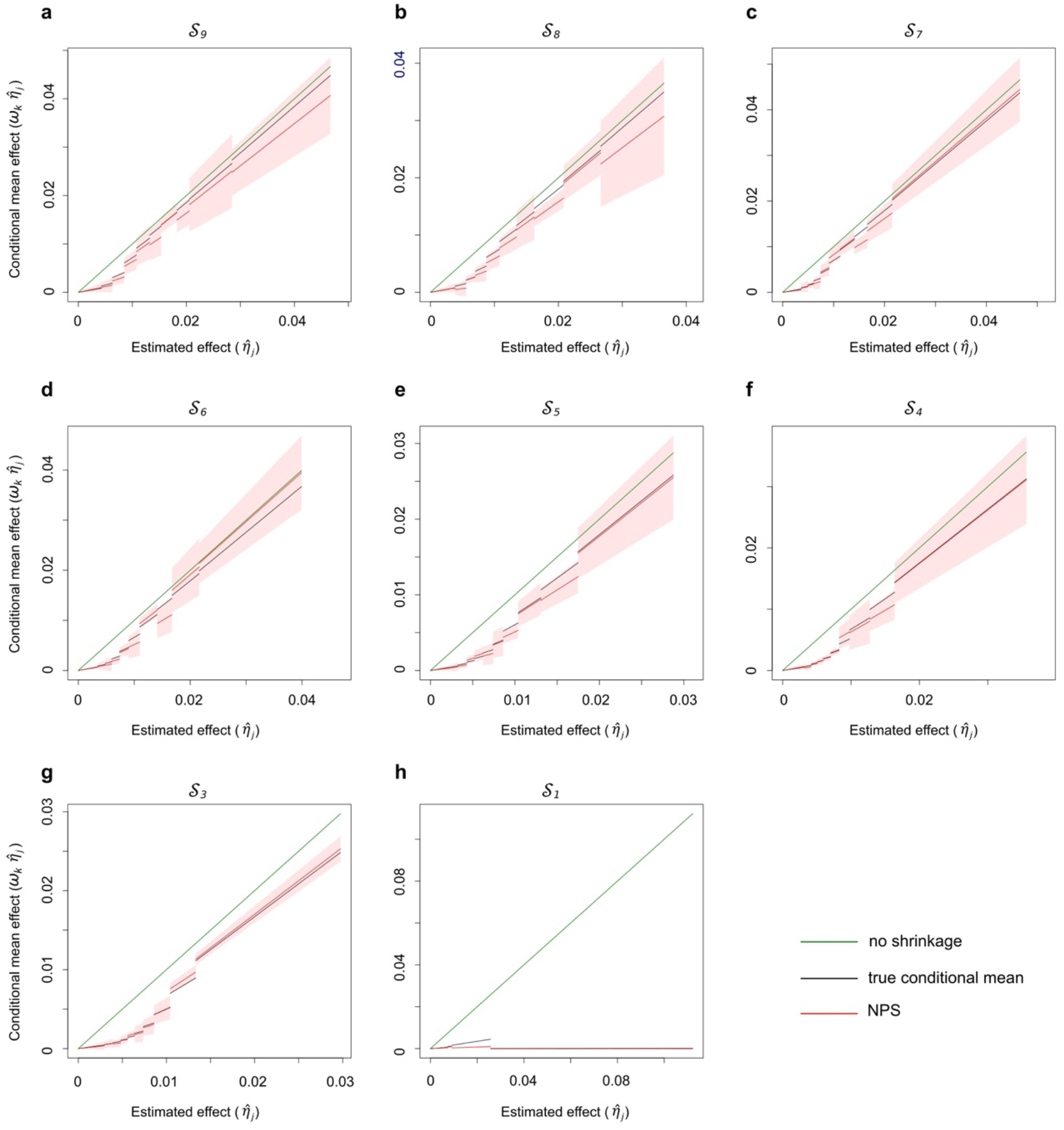
Non-parametric shrinkage (NPS) approximates the conditional mean effects: non-infinitesimal genetic architecture 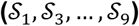. NPS shrinkage weights *ω_k_* (red line) are compared to the true conditional means (black line), which were estimated empirically from 40 simulation runs. 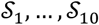 indicate the partitions of lowest to highest eigenvalues of projection. The mean NPS shrinkage weights (red line) and their 95% CIs (red shade) were estimated from 5 replicates. No shrinkage line (green) indicates *ω_k_* = 1. The number of markers *M* is 101,296. The discovery GWAS size *N* equals to *M*. The heritability *h*^2^ is 0.5. The fraction of causal SNPs is 1%. See Figure 2C-D for 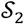 and 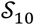, respectively.

**Figure S3.**
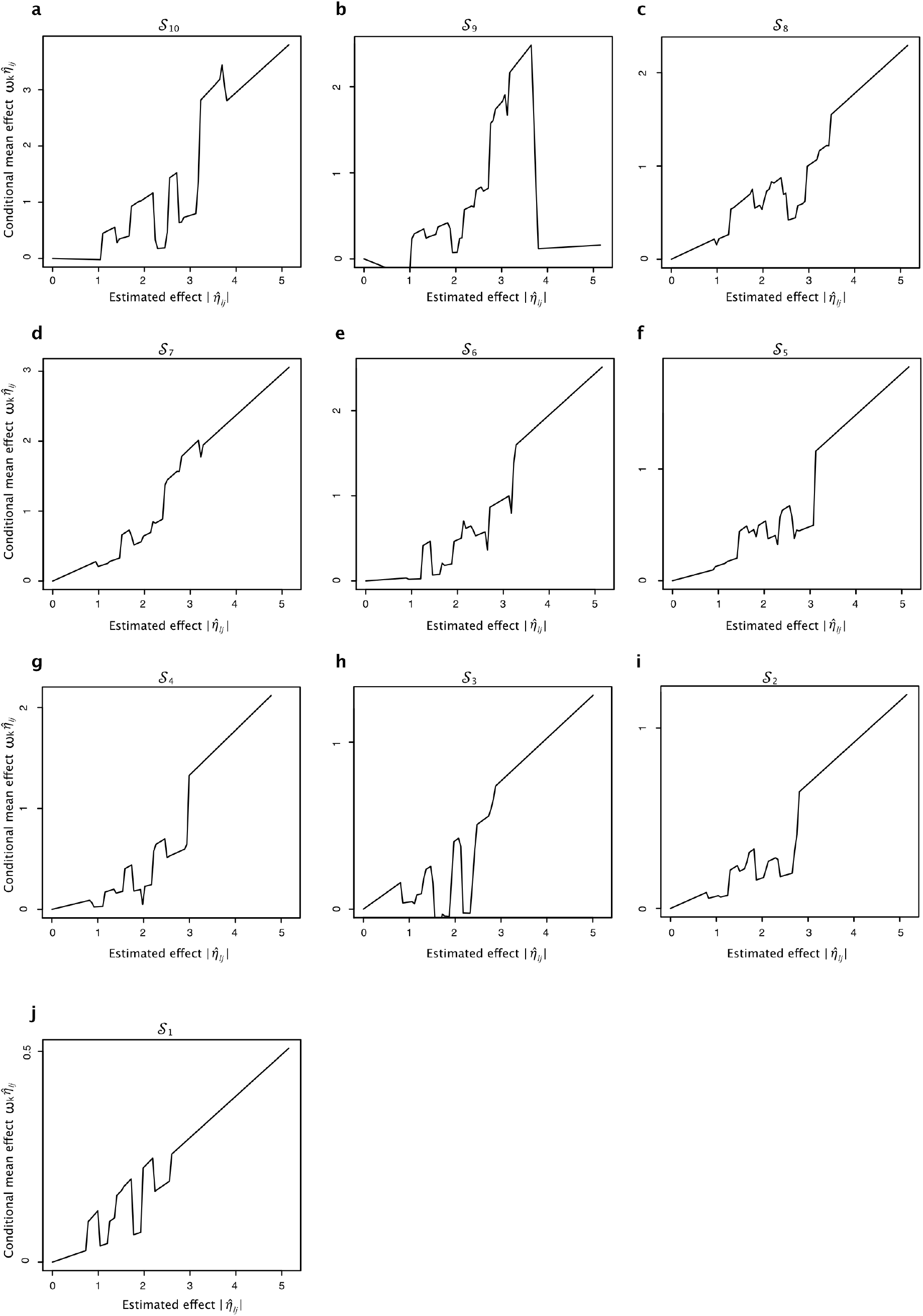
Conditional mean effects estimated by NPS in breast cancer dataset (Michailidou et al. 2017). Conditional mean effects were averaged over the four NPS runs of which windows were shifted by 0, 1,000, 2,000 and 3,000. 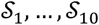 denote the partitions of lowest to highest eigenvalues of eigenlocus projection. The weights *ω_k_* were re-scaled so that the weight *ω*_0_ of genome-wide significant partition 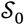 becomes 1. GWAS summary statistics are from Michailidou et al. 2017.

**Figure S4.**
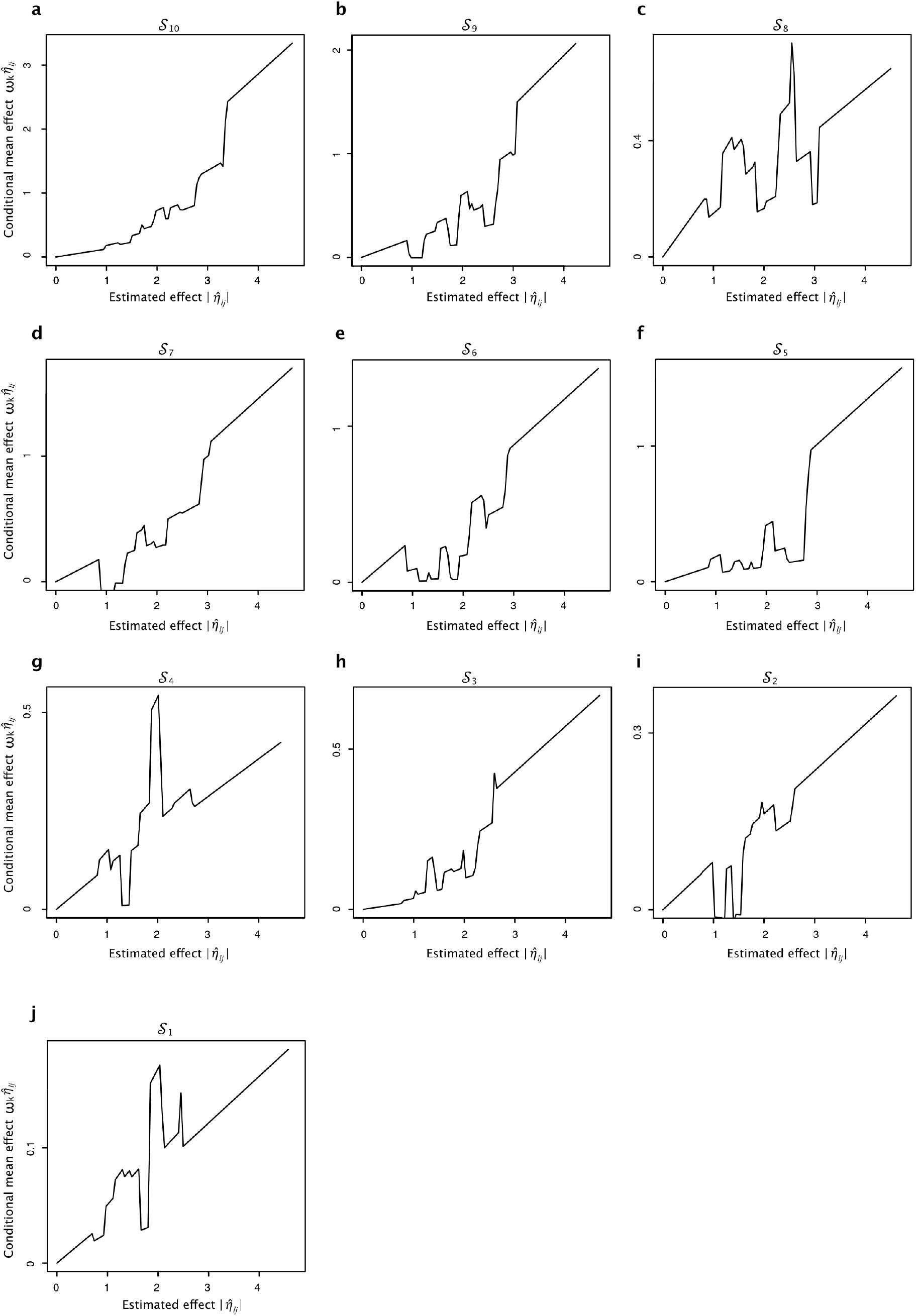
Conditional mean effects estimated by NPS in breast cancer dataset (Michailidou et al. 2015). Conditional mean effects were averaged over the four NPS runs of which windows were shifted by 0, 1,000, 2,000 and 3,000. 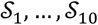 denote the partitions of lowest to highest eigenvalues of eigenlocus projection. The weights *ω_k_* were re-scaled so that the weight *ω*_0_ of genome-wide significant partition 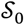 becomes 1. GWAS summary statistics are from Michailidou et al. 2015.

**Figure S5.**
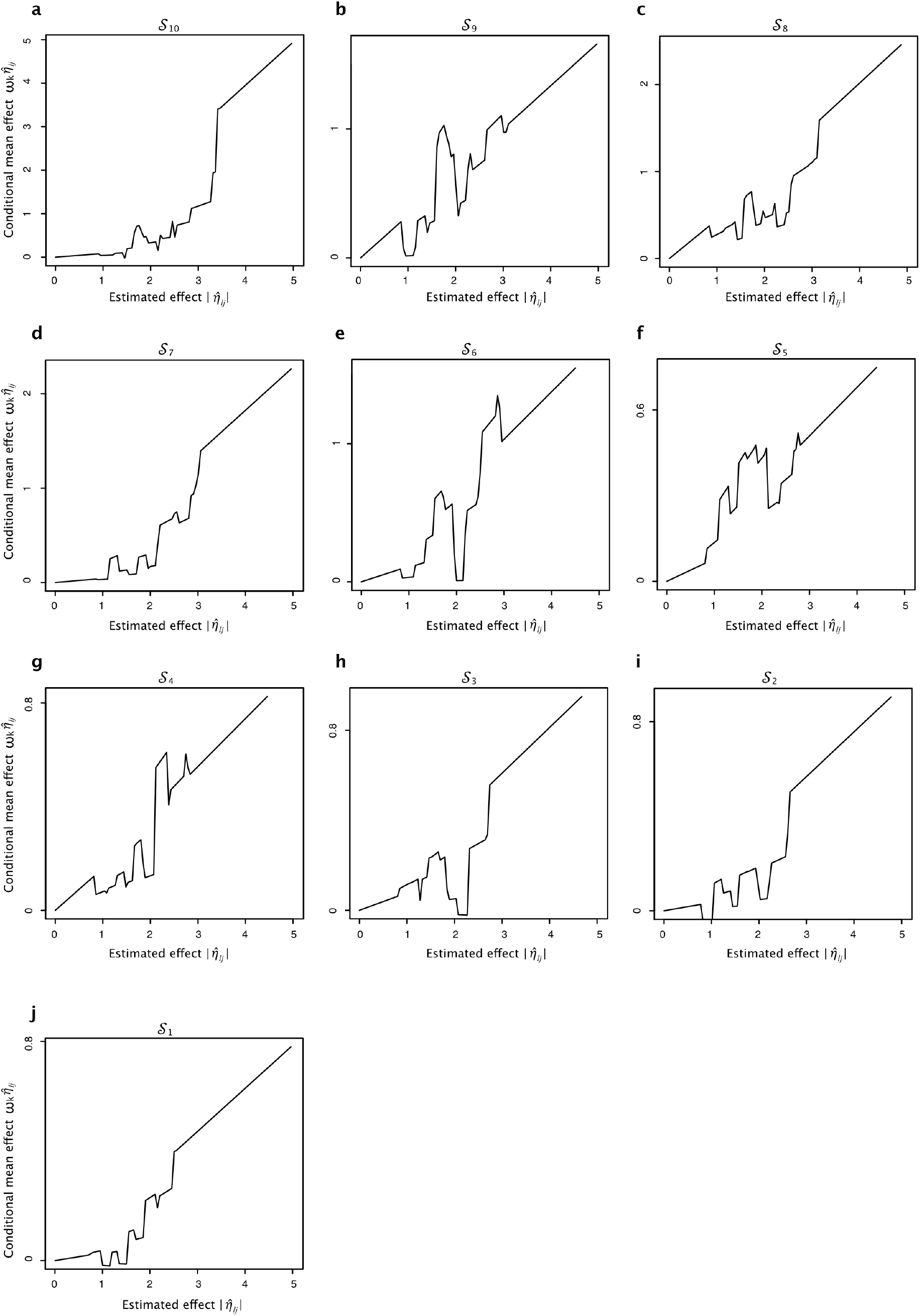
Conditional mean effects estimated by NPS in inflammatory bowel disease (IBD) dataset. Conditional mean effects were averaged over the four NPS runs of which windows were shifted by 0, 1,000, 2,000 and 3,000. 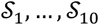 denote the partitions of lowest to highest eigenvalues of eigenlocus projection. The weights *ω_k_* were re-scaled so that the weight *ω*_0_ of genome-wide significant partition 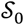 becomes 1. GWAS summary statistics are from Liu et al. 2015.

**Figure S6.**
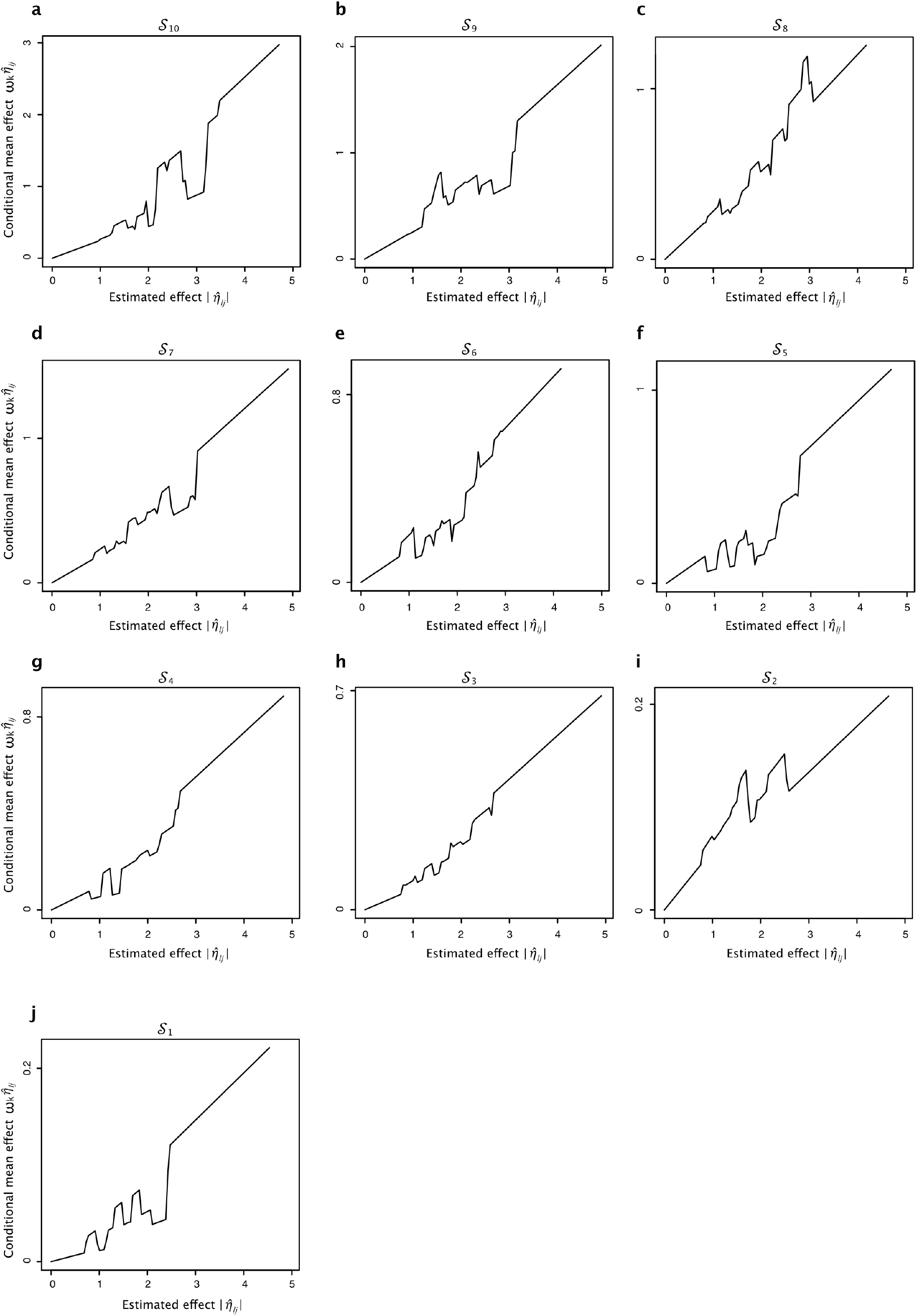
Conditional mean effects estimated by NPS in type 2 diabetes dataset. Conditional mean effects were averaged over the four NPS runs of which windows were shifted by 0, 1,000, 2,000 and 3,000. 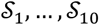 denote the partitions of lowest to highest eigenvalues of eigenlocus projection. The weights *ω_k_* were re-scaled so that the weight *ω*_0_ of genome-wide significant partition 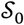 becomes 1. GWAS summary statistics are from Scott et al. 2017.

**Figure S7.**
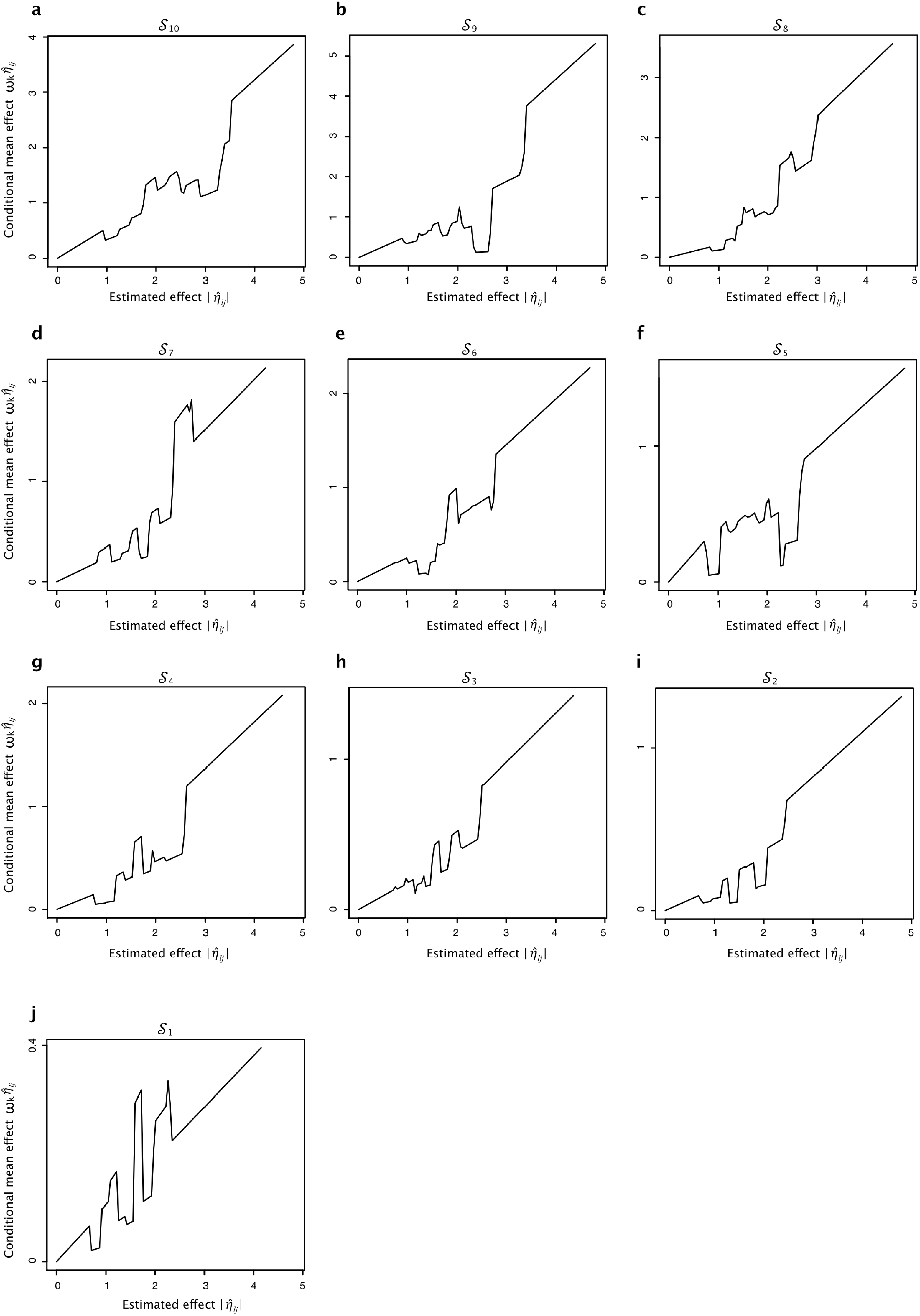
Conditional mean effects estimated by NPS in cardiovascular disease dataset. Conditional mean effects were averaged over the four NPS runs of which windows were shifted by 0, 1,000, 2,000 and 3,000. 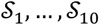 denote the partitions of lowest to highest eigenvalues of eigenlocus projection. The weights *ω_k_* were re-scaled so that the weight *ω*_0_ of genome-wide significant partition 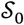 becomes 1. GWAS summary statistics are from Nelson et al. 2017.

**Table S1.**
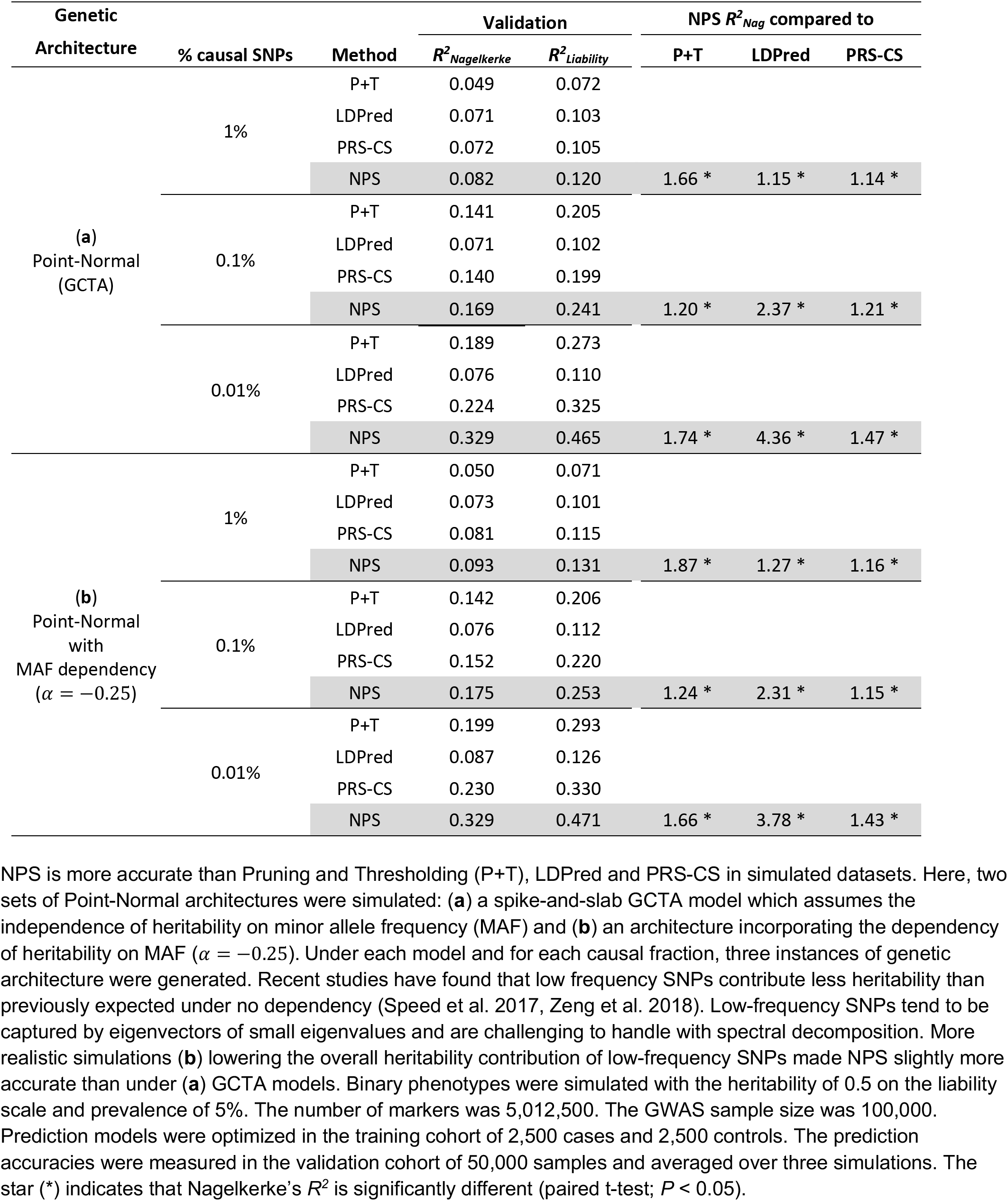
Comparison of prediction accuracy in genetic architectures simulating uniformly distributed causal SNPs.

**Table S2.** Accuracy of NPS in genetic architectures simulating the enrichment of causal SNPs within DNase I Hypersensitive Sites (DHS).

| Fraction of causal SNPs | Training | Validation |  |  |
| --- | --- | --- | --- | --- |
| | AUC | $R^2_{Nagelkerke}$ | $R^2_{Liability}$ | AUC |
| 1% | 0.746 | 0.082 | 0.126 | 0.708 |
|  | 0.737 | 0.083 | 0.125 | 0.708 |
|  | 0.725 | 0.089 | 0.118 | 0.716 |
| 0.1% | 0.800 | 0.174 | 0.249 | 0.793 |
|  | 0.811 | 0.188 | 0.262 | 0.808 |
|  | 0.802 | 0.179 | 0.261 | 0.798 |
|  | 0.810 | 0.176 | 0.254 | 0.798 |
|  | 0.810 | 0.176 | 0.250 | 0.798 |
|  | 0.813 | 0.178 | 0.259 | 0.799 |
| 0.01% | 0.891 | 0.325 | 0.463 | 0.887 |
|  | 0.894 | 0.323 | 0.462 | 0.885 |
|  | 0.887 | 0.336 | 0.463 | 0.889 |

**Table S3.**
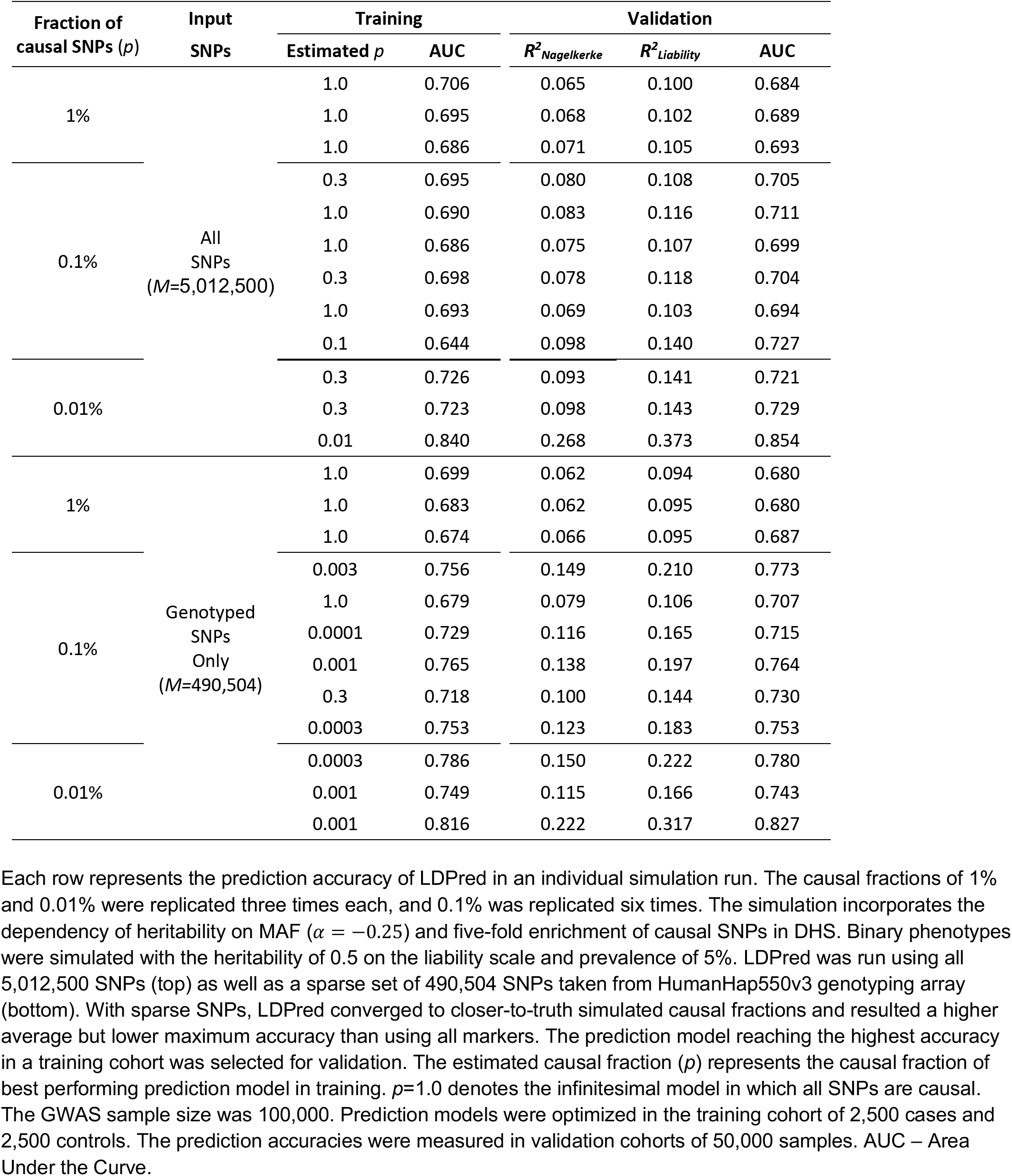
Accuracy of LDPred in genetic architectures simulating the enrichment of causal SNPs within DNase I Hypersensitive Sites (DHS).

**Table S4.** Accuracy of pruning and thresholding in genetic architectures simulating the enrichment of causal SNPs within DNase I Hypersensitive Sites (DHS).

| Fraction of causal SNPs | Training |  |  | Validation |  |  |
| --- | --- | --- | --- | --- | --- | --- |
| | P cutoff | # SNPs | AUC | $R^2_{Nagelkerke}$ | $R^2_{Liability}$ | AUC |
| 1% | 0.046 | 57,816 | 0.680 | 0.047 | 0.072 | 0.662 |
|  | 0.097 | 92,163 | 0.661 | 0.050 | 0.076 | 0.664 |
|  | 0.153 | 121,820 | 0.664 | 0.054 | 0.075 | 0.670 |
| 0.1% | 0.0001 | 2,082 | 0.783 | 0.174 | 0.244 | 0.793 |
|  | 0.00015 | 2,562 | 0.751 | 0.133 | 0.186 | 0.761 |
|  | 0.0002 | 2,765 | 0.735 | 0.119 | 0.164 | 0.747 |
|  | 0.0001 | 2,147 | 0.795 | 0.160 | 0.247 | 0.787 |
|  | 0.0001 | 2,296 | 0.736 | 0.105 | 0.163 | 0.738 |
|  | 0.00015 | 2,529 | 0.759 | 0.128 | 0.190 | 0.757 |
| 0.01% | 0.0001 | 1,662 | 0.827 | 0.209 | 0.305 | 0.823 |
|  | 0.0001 | 1,631 | 0.807 | 0.176 | 0.263 | 0.797 |
|  | 0.0001 | 1,553 | 0.833 | 0.252 | 0.352 | 0.848 |

Each row represents the prediction accuracy of pruning and thresholding (P+T) algorithm in an individual simulation run. The causal fractions of 1% and 0.01% were replicated three times each, and the causal fraction of 0.1% were replicated six times. The simulation incorporates the dependency of heritability on minor allele frequency ( $\alpha = -0.25$ ) and five-fold enrichment of causal SNPs in DHS elements. Binary phenotypes were simulated with the heritability of 0.5 on the liability scale and prevalence of 5%. The prediction model reaching the highest accuracy in a training cohort was selected for validation. The P-value cutoff of best-performing model is reported here along with the number of SNPs after pruning and thresholding. The GWAS sample size was 100,000. Prediction models were optimized in the training cohort of 2,500 cases and 2,500 controls. The prediction $R^2$ was measured in validation cohorts of 50,000 samples. AUC – Area Under the Curve.

**Table S5.** Accuracy of PRS-CS in genetic architectures simulating the enrichment of causal SNPs within DNase I Hypersensitive Sites (DHS).

| Fraction of causal SNPs | Training |  | Validation |  |  |
| --- | --- | --- | --- | --- | --- |
| | $\hat{\phi}$ | AUC | $R^2_{Nag}$ | $R^2_{Liability}$ | AUC |
| 1% | 0.01 | 0.720 | 0.072 | 0.110 | 0.693 |
|  | 0.0001 | 0.696 | 0.074 | 0.107 | 0.697 |
|  | 0.01 | 0.700 | 0.079 | 0.113 | 0.705 |
| 0.1% | 0.0001 | 0.771 | 0.157 | 0.221 | 0.780 |
|  | 0.0001 | 0.773 | 0.164 | 0.227 | 0.789 |
|  | 0.0001 | 0.769 | 0.155 | 0.224 | 0.781 |
|  | 0.0001 | 0.782 | 0.156 | 0.226 | 0.781 |
|  | 0.0001 | 0.768 | 0.148 | 0.217 | 0.777 |
|  | 0.0001 | 0.777 | 0.157 | 0.229 | 0.781 |
| 0.01% | 0.000001 | 0.835 | 0.230 | 0.332 | 0.835 |
|  | 0.000001 | 0.835 | 0.222 | 0.322 | 0.830 |
|  | 0.000001 | 0.819 | 0.232 | 0.326 | 0.833 |

Each row represents the prediction accuracy of PRS-CS in an individual simulation run. The causal fractions of 1% and 0.01% were replicated three times each, and the causal fraction of 0.1% were replicated six times. The simulation incorporates the dependency of heritability on minor allele frequency ( $\alpha = -0.25$ ) and five-fold enrichment of causal SNPs in DHS elements. Binary phenotypes were simulated with the heritability of 0.5 on the liability scale and prevalence of 5%. The prediction model reaching the highest accuracy in a training cohort was selected for validation. $\hat{\phi}$ denotes the model parameter $\phi$ of best-performing model in training. The reference LD panel was derived from a cohort sampled under the same LD structure. The GWAS sample size was 100,000. Prediction models were optimized in the training cohort of 2,500 cases and 2,500 controls. The prediction $R^2$ was measured in validation cohorts of 50,000 samples. AUC – Area Under the Curve.

**Table S6.** Accuracy of NPS applied to real GWAS summary statistics and UK Biobank datasets.

| GWAS | Training |  | Validation (UK Biobank) |  |
| --- | --- | --- | --- | --- |
|  | # Projections | AUC | AUC | Tail OR (5%) |
| Breast Cancer 2015 | 120,886 | 0.656 | 0.627 [0.62-0.64] | 2.53 [2.3-2.8] |
| Breast Cancer 2017 | 124,061 | 0.678 | 0.654 [0.65-0.66] | 3.01 [2.7-3.3] |
| IBD | 110,157 | 0.686 | 0.659 [0.65-0.67] | 3.60 [3.2-4.0] |
| Type 2 Diabetes | 139,106 | 0.697 | 0.686 [0.68-0.69] | 3.81 [3.6-4.1] |
| CAD | 105,162 | 0.778 | 0.738 [0.72-0.76] | 5.21 [4.3-6.2] |

**Table S7.** Accuracy of LDPred applied to real GWAS summary statistics and UK Biobank datasets.

| GWAS | Training |  |  | Validation (UK Biobank) |  |
| --- | --- | --- | --- | --- | --- |
|  | # SNPs | Estimated causal fraction | AUC | AUC | Tail OR (5%) |
| Breast Cancer 2015 | 3,417,759 | 0.01 | **0.630** | 0.618 [0.61-0.63] | 2.42 [2.2-2.7] |
| Breast Cancer 2017 | 3,478,993 | 0.1 | **0.621** | 0.615 [0.61-0.62] | 2.33 [2.1-2.6] |
| IBD | 3,396,783 | 0.03 | **0.640** | 0.641 [0.63-0.65] | 2.77 [2.4-3.1] |
| Type 2 Diabetes | 3,451,818 | 0.01 | **0.680** | 0.679 [0.67-0.68] | 3.51 [3.3-3.8] |
| CAD | 3,405,299 | 0.003 | 0.753 | 0.738 [0.72-0.76] | 5.17 [4.3-6.1] |
| Breast Cancer 2015 | 351,917 | 0.3 | 0.605 | 0.597 [0.59-0.61] | 2.25 [2.0-2.5] |
| Breast Cancer 2017 | 353,627 | 1.0 | 0.606 | 0.604 [0.60-0.61] | 2.03 [1.8-2.3] |
| IBD | 353,325 | 1.0 | 0.618 | 0.622 [0.61-0.63] | 2.76 [2.4-3.1] |
| Type 2 Diabetes | 354,110 | 0.1 | 0.679 | 0.680 [0.67-0.69] | 3.63 [3.4-3.9] |
| CAD | 329,644 | 0.03 | **0.757** | 0.742 [0.72-0.76] | 5.65 [4.7-6.7] |

**Table S8.**
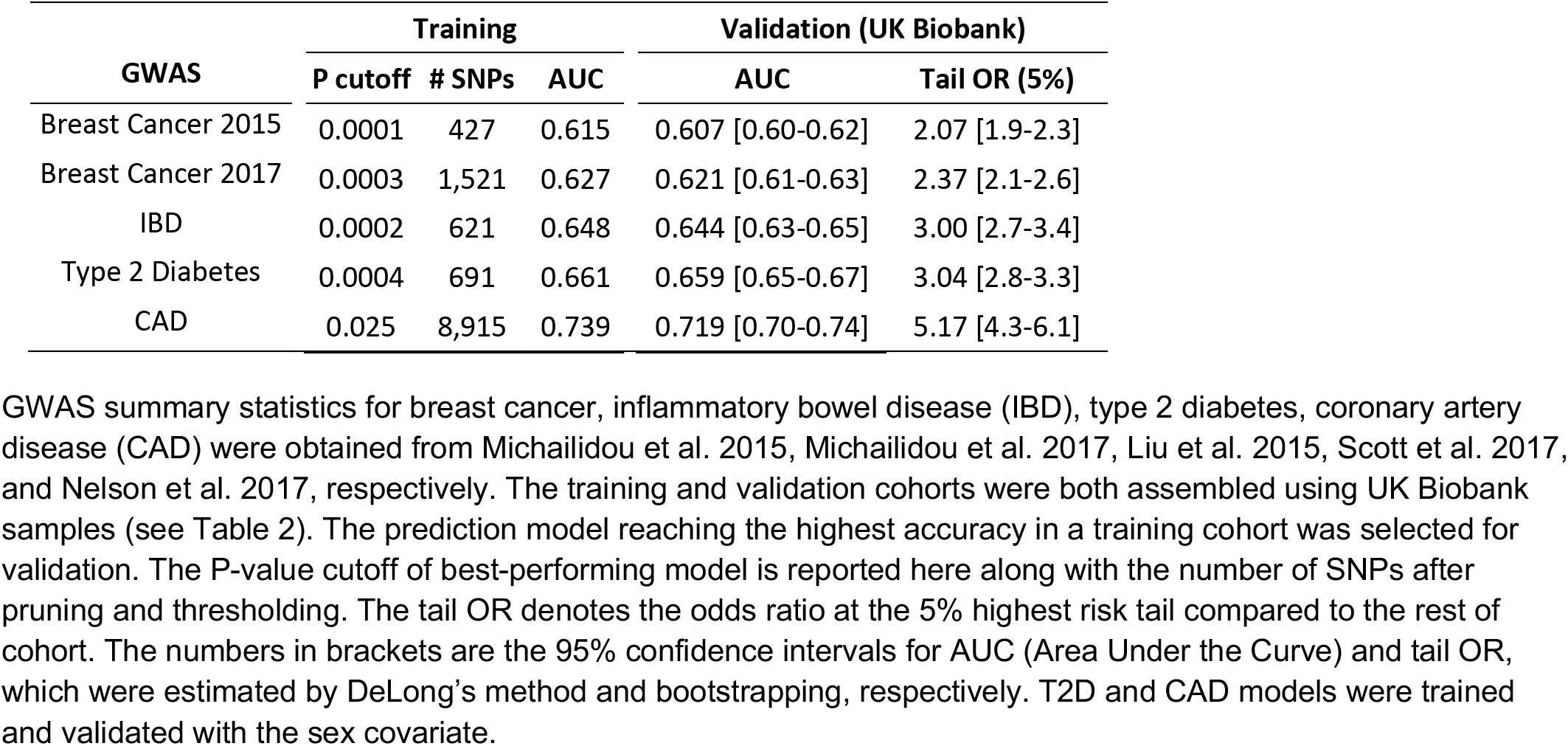
Accuracy of pruning and thresholding applied to real GWAS summary statistics and UK Biobank datasets.

| GWAS | Training |  |  | Validation (UK Biobank) |  |
| --- | --- | --- | --- | --- | --- |
|  | P cutoff | # SNPs | AUC | AUC | Tail OR (5%) |
| Breast Cancer 2015 | 0.0001 | 427 | 0.615 | 0.607 [0.60-0.62] | 2.07 [1.9-2.3] |
| Breast Cancer 2017 | 0.0003 | 1,521 | 0.627 | 0.621 [0.61-0.63] | 2.37 [2.1-2.6] |
| IBD | 0.0002 | 621 | 0.648 | 0.644 [0.63-0.65] | 3.00 [2.7-3.4] |
| Type 2 Diabetes | 0.0004 | 691 | 0.661 | 0.659 [0.65-0.67] | 3.04 [2.8-3.3] |
| CAD | 0.025 | 8,915 | 0.739 | 0.719 [0.70-0.74] | 5.17 [4.3-6.1] |

**Table S9.** Accuracy of PRS-CS applied to real GWAS summary statistics and UK Biobank datasets.

| GWAS | Training |  |  | Validation (UK Biobank) |  |
| --- | --- | --- | --- | --- | --- |
| | # SNPs | $\hat{\phi}$ | AUC | AUC | Tail OR (5%) |
| Breast Cancer 2015 | 712,303 | 0.0001 | 0.635 | 0.626 [0.62-0.63] | 2.60 [2.3-2.9] |
| Breast Cancer 2017 | 711,549 | 0.0001 | 0.657 | 0.651 [0.64-0.66] | 2.96 [2.7-3.3] |
| IBD | 707,371 | 0.0001 | 0.665 | 0.668 [0.66-0.68] | 3.67 [3.3-4.1] |
| Type 2 Diabetes | 715,952 | 0.0001 | 0.686 | 0.688 [0.68-0.69] | 3.99 [3.7-4.3] |
| CAD | 708,976 | 0.0001 | 0.763 | 0.739 [0.72-0.76] | 4.92 [4.1-5.8] |

**Table S10.** Accuracy of NPS in independent validation cohorts.

| GWAS | Training |  | Validation (Partners Biobank) |  |
| --- | --- | --- | --- | --- |
|  | # Projections | AUC | AUC | Tail OR (5%) |
| Breast Cancer 2017 | 124,061 | 0.678 | 0.624 [0.60-0.64] | 2.32 [1.7-3.0] |
| IBD | 110,157 | 0.686 | 0.686 [0.67-0.70] | 4.32 [3.5-5.2] |
| Type 2 Diabetes | 139,106 | 0.697 | 0.647 [0.63-0.66] | 2.97 [2.6-3.5] |
| CAD | 105,162 | 0.778 | 0.615 [0.58-0.65] | 4.10 [2.8-5.8] |

**Table S11.**
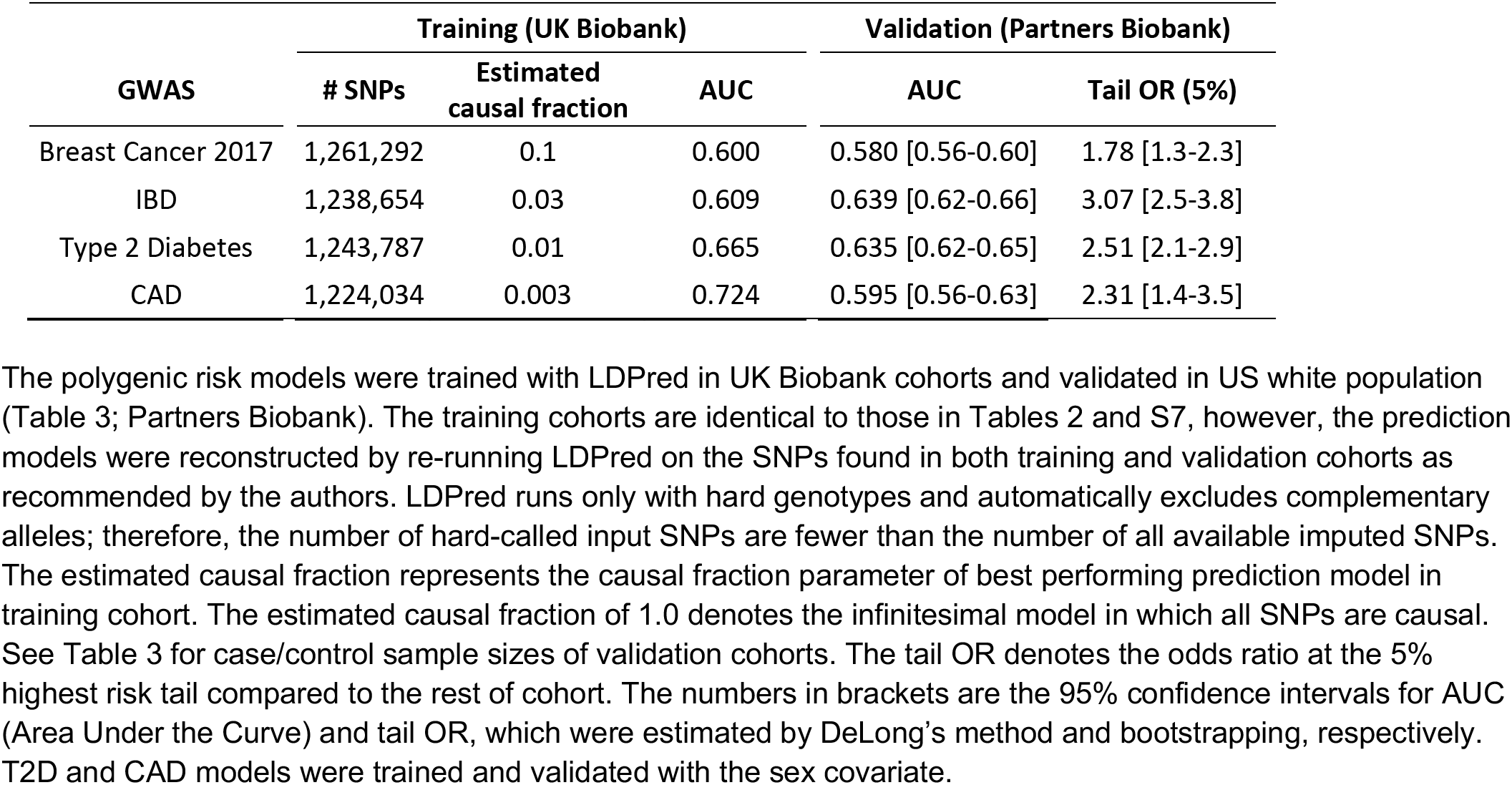
Accuracy of LDPred in independent validation cohorts.

| GWAS | Training (UK Biobank) |  |  | Validation (Partners Biobank) |  |
| --- | --- | --- | --- | --- | --- |
|  | # SNPs | Estimated causal fraction | AUC | AUC | Tail OR (5%) |
| Breast Cancer 2017 | 1,261,292 | 0.1 | 0.600 | 0.580 [0.56-0.60] | 1.78 [1.3-2.3] |
| IBD | 1,238,654 | 0.03 | 0.609 | 0.639 [0.62-0.66] | 3.07 [2.5-3.8] |
| Type 2 Diabetes | 1,243,787 | 0.01 | 0.665 | 0.635 [0.62-0.65] | 2.51 [2.1-2.9] |
| CAD | 1,224,034 | 0.003 | 0.724 | 0.595 [0.56-0.63] | 2.31 [1.4-3.5] |

**Table S12.** Accuracy of pruning and thresholding in independent validation cohorts.

| GWAS | Training (UK Biobank) |  |  | Validation (Partners Biobank) |  |
| --- | --- | --- | --- | --- | --- |
|  | P cutoff | # SNPs | AUC | AUC | Tail OR (5%) |
| Breast Cancer 2017 | 0.00035 | 801 | 0.613 | 0.589 [0.57-0.61] | 1.56 [1.2-2.1] |
| IBD | 0.0002 | 331 | 0.629 | 0.659 [0.64-0.68] | 3.57 [2.9-4.4] |
| Type 2 Diabetes | 0.0001 | 165 | 0.656 | 0.623 [0.61-0.64] | 2.10 [1.8-2.5] |
| CAD | 0.15 | 15,908 | 0.739 | 0.611 [0.58-0.65] | 2.72 [1.8-3.9] |

**Table S13.**
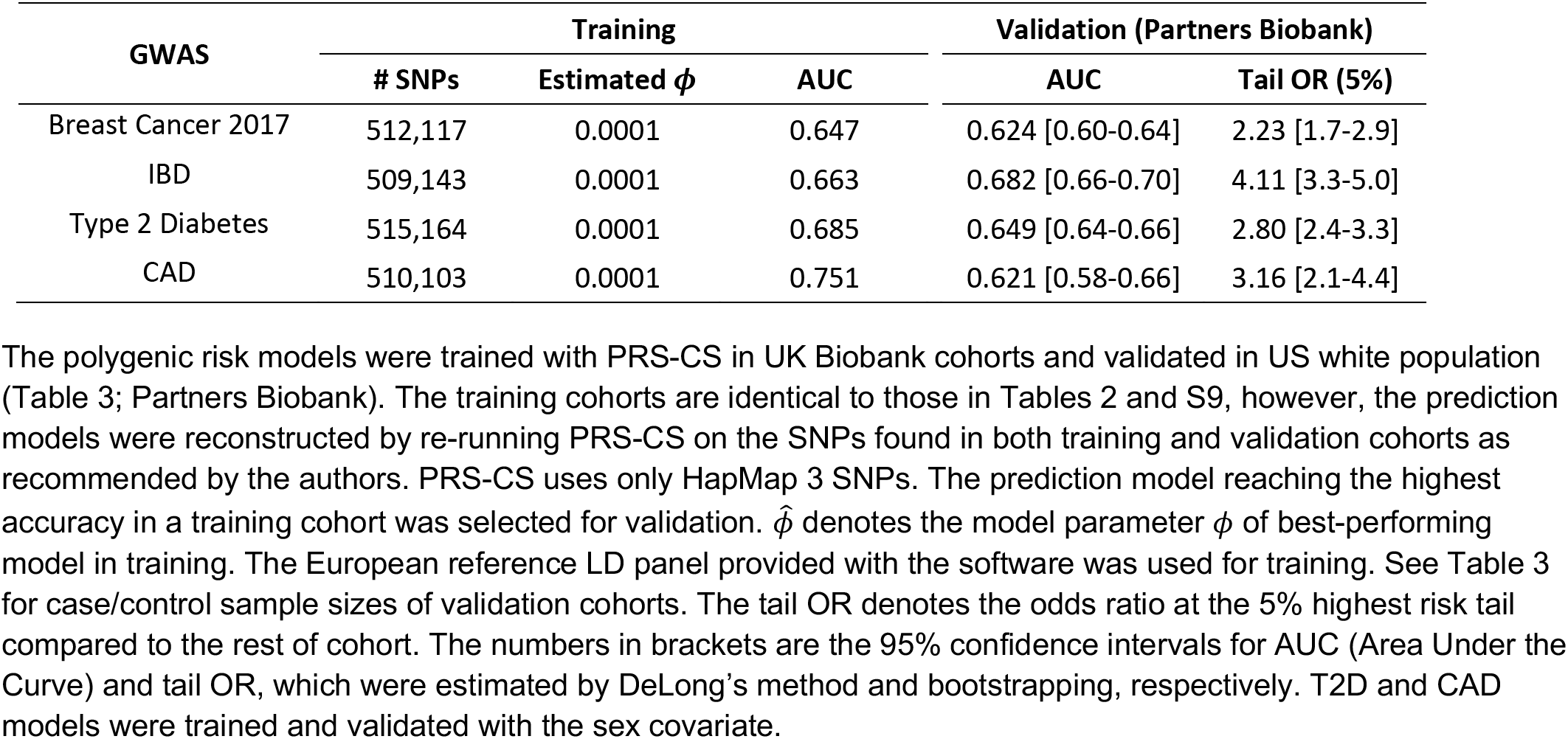
Accuracy of PRS-CS in independent validation cohorts.

